# The geometry of context-dependent biased decisions during learning

**DOI:** 10.64898/2026.01.14.699497

**Authors:** Ramon Nogueira, Saleh Esteki, Stefano Fusi, Roozbeh Kiani

## Abstract

Adaptive behavior requires inferring latent context and rapidly adjusting decisions in response to changing environmental contingencies. We investigated how reward context is learned, represented, and updated during decision making. We recorded large populations of neurons in lateral prefrontal cortex while macaque monkeys learned a direction-discrimination task in which reward contingencies alternated unpredictably between favoring leftward and rightward choices. Once trained, monkeys inferred context switches from a single unexpected outcome, immediately adjusting both choice bias and reaction times—hallmarks of model-based inference. Early in learning, however, adaptation unfolded gradually across multiple trials. Neural population analyses revealed that reward context was encoded through systematic shifts in the geometry of neural representations. Accumulated sensory evidence (decision variable) and choice were organized along curvilinear decision manifolds, which were displaced across contexts primarily along the decision-variable axis. This geometry naturally implemented context-dependent biases: a fixed linear readout generated different choice tendencies across contexts without remapping. Longitudinal recordings further showed that, with learning, these representational transitions between manifolds became faster, mirroring the emergence of one-trial behavioral generalization. Recurrent neural networks trained on the same task reproduced both the behavioral signatures and the context-dependent geometric shifts. Together, these findings identify a mechanism by which prefrontal circuits support hierarchical inference: reward context is encoded as structured shifts in representational geometry, enabling rapid generalization and flexible control of decision policies.

## Introduction

Decision making is inherently hierarchical.^1, 2^ Decision makers must not only evaluate immediate sensory evidence but also monitor broader environmental contingencies and adjust behavior accordingly.^1, 3–6^ This flexibility—switching strategies when conditions change—relies on frontoparietal and subcortical circuits that integrate sensory evidence with contextual signals such as sensory-motor mappings, expected rewards, priors, and task rules.^2, 7–17^ A central component of this computation is expected reward,^18, 19^ which biases choice behavior: individuals favor options with greater expected value and commit to them more quickly.^20–22^ These behavioral signatures have been formalized in normative and mechanistic models of decision making and traced to the dynamics of single neurons in association cortices.^11, 21–26^ Yet, how these signals are coordinated across levels of representation to support hierarchical inference remains poorly understood.

We focus here on two critical questions about the effect of expected rewards on decision making. First, how are abstract reward contexts—regularities that shape the mapping from evidence to action—learned and represented in the brain? Second, how are sudden changes in reward context inferred and incorporated into ongoing computations to alter choices and reaction times? To address these questions, we chronically recorded from large populations of neurons in the prearcuate gyrus (area 8Ar) of the lateral prefrontal cortex (lPFC) while monkeys learned and performed an alternating-context direction discrimination task. Monkeys reported the direction of a random dot kinematogram with a saccadic eye movement^27, 28^ (Fig. 1a), while reward contingencies alternated between favoring leftward and rightward choices. Critically, context switches were uncued; they occurred at random intervals (200 trials on average) and had to be inferred from feedback (Fig. 1b). Task difficulty was varied trial by trial by changing the percentage of coherently moving dots (motion strength), allowing us to quantify the interaction of reward context with the decision-making process.^29^ Once monkeys had learned the task, they rapidly adapted after a single unexpected feedback, shifting both their choice bias and reaction times toward the newly favored option—a behavioral signature of context inference for normative decisions. Early in training, however, adaptation was gradual and spanned across several trials. The behavioral transition from novice to expert was mirrored by systematic changes in lPFC neural activity.

**Figure 1:**
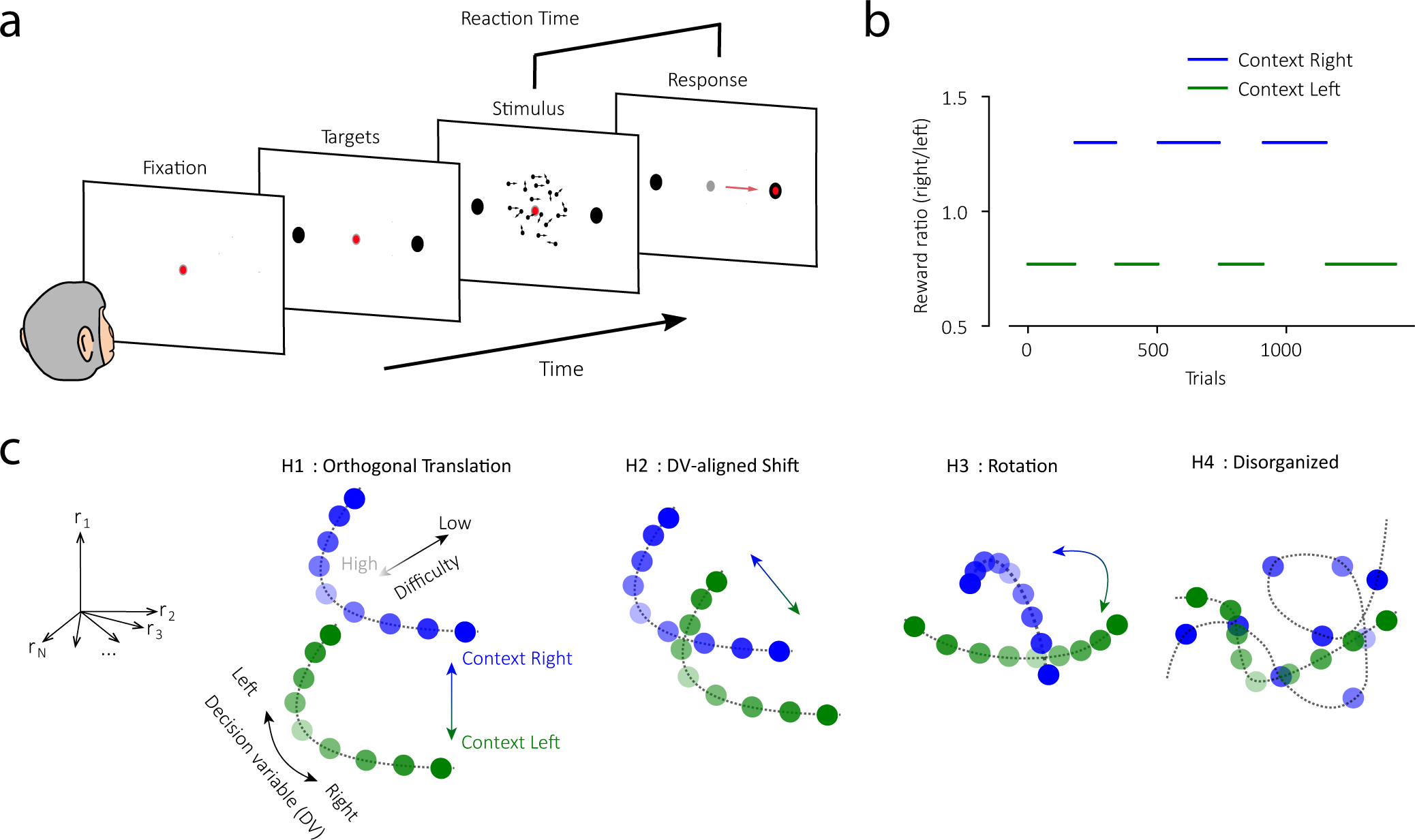
The alternating-context direction discrimination task and differential hypotheses about representational geometries. (**a**) On each trial, monkeys viewed a random dot motion stimulus and reported the net motion direction (left or right) with a saccade to one of two peripheral targets. We controlled stimulus difficulty by varying the percentage of coherently moving dots (0%-51.2%). Monkeys controlled viewing duration and initiated the saccade when ready (red dot shows gaze position in different task epochs, and red arrow indicates saccade direction). Reaction time was defined as the interval between stimulus onset and saccade initiation. (**b**) Example session showing alternating reward contexts. In Context Right (blue), correct rightward choices yielded larger rewards than leftward choices. The reward asymmetry was reversed in Context Left (green). Contexts lasted for many trials and switched unpredictably without explicit cues. (**c**) Schematic representational geometries. In H1-H3, lPFC population activity is organized along axes for the decision variable (DV; accumulated sensory evidence supporting the choice) and stimulus difficulty (motion coherence; light vs. dark shading) in each reward context, but changes systematically across contexts (green vs. blue). H1: reward context translates the decision manifold in a direction orthogonal to the DV and stimulus difficulty. H2: reward context shifts the manifold along the DV axis, biasing a fixed readout. H3: context rotates the manifold, requiring context-aware readout. H4: representations across contexts are unrelated and high-dimensional.

Analysis of population responses in lPFC revealed that context learning reshaped the geometry of neural representations. While we also analyzed single-unit responses, we focus here on population activity patterns and changes in representational geometry, which offer an intuitive and computationally informative perspective on neural data. Past work has shown that representational geometry reveals interpretable computational structures^30–32^ that are conserved across individuals.^33, 34^ Low-dimensional geometries, in which task variables are represented in approximately orthogonal subspaces (disentangled representations), have been observed in the hippocampus and prefrontal cortex of non-human^35, 36^ and human primates,^37^ in inferotemporal cortex of monkeys,^38^ in the mouse somatosensory cortex^39^ and hippocampus,^40^ and in decision-making tasks.^13, 41, 42^ These representations are typically the result of an abstraction process and allow for simple forms of out-of-distribution generalization. Other studies, however, have revealed high-dimensional representations in macaque prefrontal cortex, with dimensionality correlating with performance in complex tasks.^43, 44^

In our task, neural activity for each combination of motion strength and choice formed curvilinear population response manifolds, where activity patterns associated with similar task conditions were closer to each other.^41, 45^ We tested different hypotheses about how reward context altered these manifolds (Fig. 1). The two manifolds that correspond to different contexts could have the same shape, but shifted in a direction orthogonal to the curvilinear manifolds (H1). Under this hypothesis, a single linear readout for choice would readily work for both contexts. However, a simple readout represented by a plane orthogonal to the manifolds would not explain changes in the behavioral bias. Alternatively, the two manifolds could be shifted along the axis that represents the monkey’s decision variable (H2). With a linear readout, this shift yields a context-dependent behavioral bias. A third possibility is that the two manifolds could be translated and rotated with respect to each other (H3), as has been shown for rule changes in the parietal cortex.^41^ It is also possible that a change of reward context alters the manifolds in an unstructured way in the high-dimensional activity space (H4). H3 and H4 preserve context information but require context-aware readouts^41^ and lack abstraction of task structure.^35^

We show that the manifolds corresponding to different contexts were separable and mainly shifted along the decision-variable axis, supporting the observed behavioral biases. This geometry allowed a fixed linear readout to capture context-dependent biases. Simulations with recurrent neural networks (RNNs) trained on the same task reproduced these representational shifts, supporting their computational relevance. Moreover, longitudinal recordings showed that lPFC representations evolved in parallel with behavioral learning, shifting more rapidly as monkeys acquired the ability to infer context after a single unexpected reward. Together, these findings reveal a representational mechanism by which prefrontal circuits support hierarchical decisions: reward contexts that bias choices are encoded as structured shifts in population geometry, enabling downstream circuits to generalize, infer, and adapt behavior with remarkable speed.

## Results

Monkeys performed the task well. The probability of choosing the right target increased monotonically with evidence towards rightward motion, and higher motion coherence elicited faster responses, consistent with integration of sensory evidence to a decision bound.^46–48^ Importantly, monkeys also adapted their choices to exploit the context-dependent reward imbalance: they chose the higher-paying target more frequently (psychometric shifts Δ*β*_0_ = 8.1 ± 0.6, Eq. 1, *P <* 10^−8^, two-sided paired t-test) and more quickly in each context (mean *RT* change between low- and high-paying choices, 56.7 ± 4.2 ms, *P <* 10^−8^, one-sided z-test), evidenced by systematic shifts in psychometric and chronometric functions (Fig. 2a,b). This pattern of behavior enhances cumulative reward in environments with reward asymmetry and is well explained by a dynamic bias signal that pushes the decision variable toward the decision bound for the higher-paying target.^11, 24^

**Figure 2:**
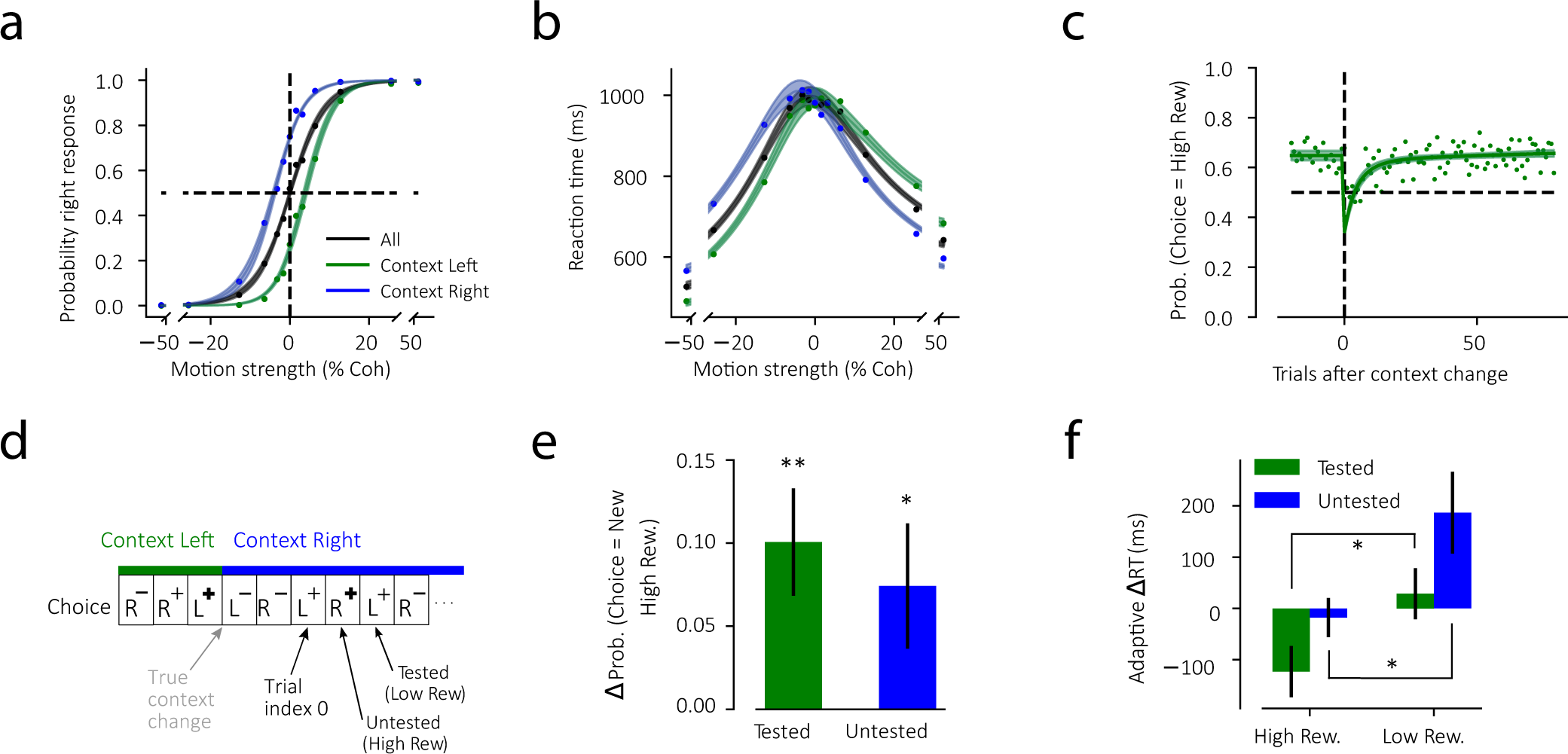
Monkeys bias their choices toward the most rewarded target and infer context based on a model of the task. (**a**) Psychometric curves: Probability of rightward choices as a function of motion strength for all trials (black), context left (green), and context right (blue). Positive and negative motion strengths indicate rightward and leftward motion directions, respectively. Choice probability was shifted toward the more rewarded target in each context. (**b**) Chronometric curves: Reaction time as a function of motion strength. Reaction times were shorter when motion direction was congruent with the context bias. (**c**) Probability of choosing the higher-reward target after an uncued context switch. Trial index 0 indicates the first rewarded trial after the switch. The experienced change of reward size on this trial is the monkey’s first piece of evidence for a context change. (**d**) Definition of tested and untested inference trials: On the tested trial, the monkey repeated the same choice as trial index 0, knowing the expected reward. On the untested trial, the monkey chose the opposite target and could know about the change of expected reward only with a model of the task. R and L indicate right and left choices, negative sign (−) indicates unrewarded trials, and thin and thick positive signs (+ and **+**) indicate small and large rewards. (**e**) Change in the probability of choosing the new higher-reward target after a context switch, relative to a pre-switch baseline, shown separately for tested (green) and untested (blue) trials. (**f**) Change in reaction time after a switch, relative to the pre-switch baseline, for choices toward the higher- and lower-reward targets. Data are pooled across the two monkeys, and error bars correspond to s.e.m. across sessions. See Supplementary Fig. S1 for individual monkeys. ∗ = *P <* 0.05, ∗∗ = *P <* 0.01

On each recording session, multiple uncued context changes occurred. Well-trained monkeys rapidly adjusted their choice biases after a context switch (Fig. 2c). The first rewarded trial after a switch provided the first piece of evidence that the context had changed. We define this first correct, post-switch trial as the subjective context switch (trial index 0), and the choice made on it as the tested target. In our task, a change in reward context alters the expected value of both choices. If monkeys had acquired a model of the task, they could generalize from the reward experienced on the tested target to the expected reward for the untested one. For example, if rewards had previously favored the leftward choice (large reward left, small reward right), and on trial 0 a rightward choice yielded a large reward, then an agent with a task model would infer that rewards for leftward choices have now become small. By contrast, an agent that simply updates reward expectations independently for each target would continue to expect a large reward for the leftward choice until it directly experienced otherwise.

We tested these alternatives by examining both choices and reaction times on trials immediately following trial 0 (Fig. 2d–f). If behavior was based on a task model, adjustments should generalize to the untested target: monkeys should increase the probability of choosing the new higher-reward option and simultaneously slow down responses to the newly devalued option, even when that option had not yet been directly experienced in the new context. In contrast, if monkeys updated each choice option independently, such generalization would not be observed. Consistent with task inference, monkeys showed immediate generalization. A single unexpected reward outcome was sufficient to bias subsequent choices toward the new higher-reward target, and this adjustment was evident for both tested and untested motion directions (Fig.2e; tested *P* = 3.2 × 10^−3^, untested *P* = 3.3 × 10^−2^, one-sided t-test). Reaction times provided a complementary analog measure. After trial 0, monkeys responded more quickly to the newly favored target and more slowly to the newly disfavored target—even when the disfavored option had not yet been tested (Fig.2f; tested *P* = 0.023, untested *P* = 0.011, one-sided t-test). These results indicate that monkeys inferred the new reward context from a model of the task, rather than relying only on direct reward experience for each choice.

### Context shifts neural representations in lPFC

While monkeys performed the task, we recorded from populations of neurons in the prearcuate region (area 8Ar), where neural responses represent integration of sensory evidence, priors, and decision rules to guide saccadic eye movements.^13, 49–51^ Among recorded single and multi-units, 52% were selective to choice, 31% to reward context, and 18% to both (Fig. 3a). The context-selective units often persistently represented the reward context throughout the block, including both the trial and inter-trial intervals, and switched their activity level following a change of reward context. In contrast, choice-selective units exhibited progressively divergent responses for the two choices, starting ∼200 ms after stimulus onset.

**Figure 3:**
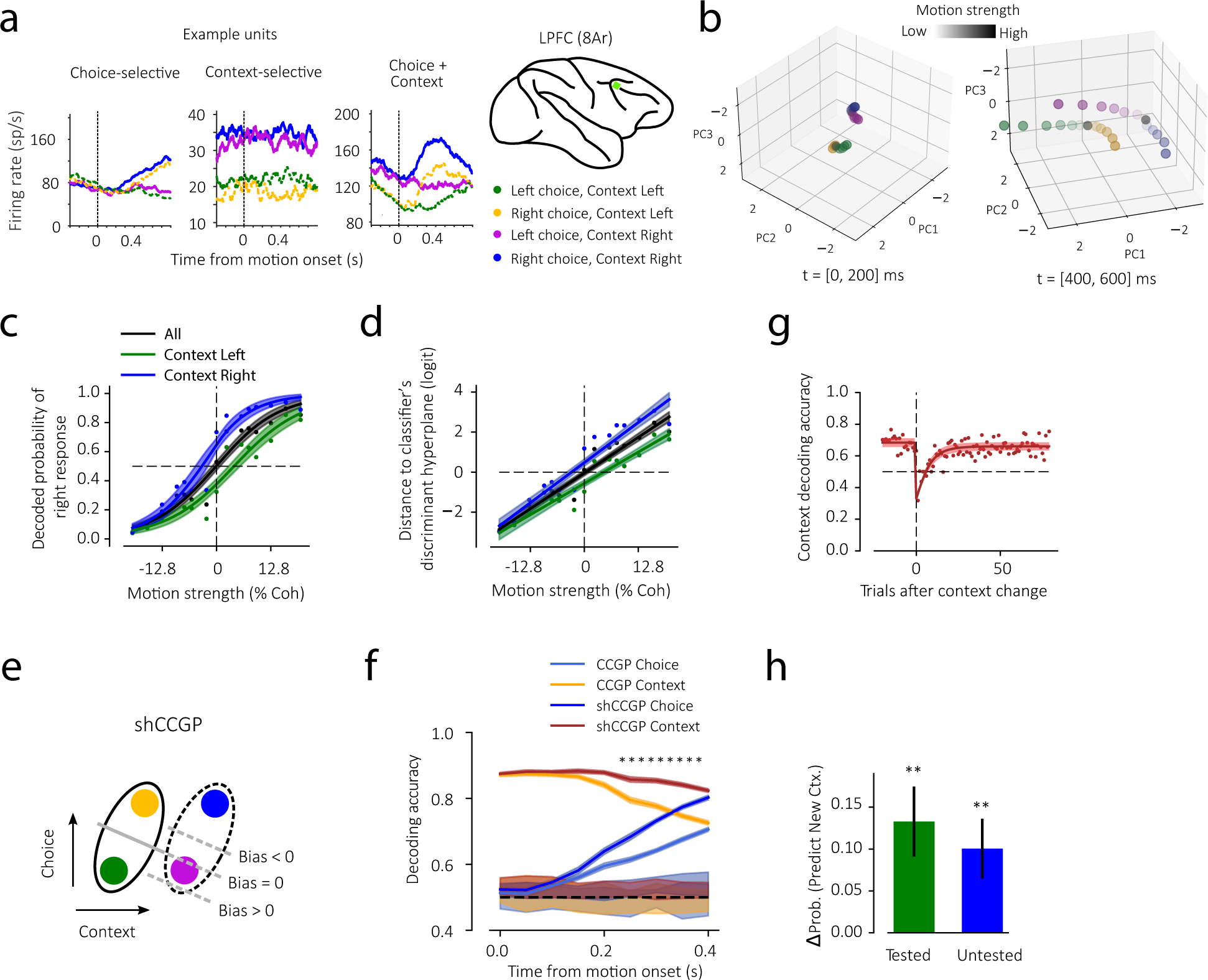
The geometry of lPFC representations supports context-dependent bias and context inference. **(a)** PSTHs of three example lPFC units selective for choice (left), reward context (middle), or both (right). (**b**) Population activity patterns projected onto the first three principal components. Early (0-200 ms after motion onset, left): activity patterns for different motion strengths are indistinguishable within a context. Late (400-600 ms, right): activity patterns spread along curvilinear manifolds reflecting motion strength and choice, with manifolds shifted along the decision variable axis across contexts. (**c**) Neurometric curves: Probability of a logistic classifier predicting a rightward choice based on the pattern of lPFC population activity (0-600 ms after motion onset) as a function of motion strength for all trials (black), context left (green), or context right (blue). Shading indicates standard deviation across different instances of pseudopopulation activity patterns (see Methods). (**d**) Distance of lPFC activity to the classifier’s discriminant hyperplane showing context-dependent shifts. (**e**) Schematic of shifted Cross-Condition Generalization Performance (shCCGP): classifier trained on one set of conditions and tested on the complementary set, with a bias term adjusted to account for manifold shift. For instance, a choice classifier (solid line) trained to discriminate right vs. left choices on context left trials (green vs. orange; solid ellipse) could be tested on context right trials (magenta vs. blue; dashed ellipse), while adjusting a bias term to produce the highest generalization performance (the top dashed line maximizes performance). (**f**) Comparison of standard CCGP (pale) and shCCGP (dark) for choice (blue) and context (brown) during the stimulus viewing epoch. Asterisks indicate when shCCGP significantly exceeds CCGP for both choice and context (non-parametric permutation test, *P <* 10^−2^). Shaded region around CCGP and shCCGP shows standard deviation across samples of pseudopopulation activity. Shaded regions around chance (DP = 0.5) correspond to the 2.5th to 97.5th percentiles of the null distribution. (**g**) Decoding accuracy of single-session lPFC population activity for reward context across trials after an uncued context switch. Trial index 0 corresponds to the first rewarded trial after a context change. (h) Change in the probability of classifying lPFC population activity as a new context after context switches. Results are split between trials that matched the choice of the first rewarded trial after a context switch (tested; green) and those that did not match (untested; blue) (see Methods). See Supplementary Fig. S4 for the results of individual monkeys. Data are pooled across the two monkeys, and error bars in g,h correspond to s.e.m. across sessions.

To visualize the geometry of these neural representations, we removed trial-to-trial noise using an encoding model (see Methods) and projected the denoised population activity onto the top three principal components (Fig. 3b; 84% variance explained). Each point in this space represented one experimental condition. We discretized motion strengths (3 leftward, 3 rightward, and zero coherence) for each reward context, yielding 14 conditions. Shortly after motion onset, the seven motion strengths within each context were tightly clustered and indistinguishable, but the two contexts remained well separated—consistent with the persistent activity of context-selective neurons across trials. As the monkey integrated sensory evidence in the trial, the points associated with different motion strengths spread along distinct curvilinear manifolds for each context. Crucially, these manifolds were systematically shifted relative to one another along the decision variable axis (Fig. 1c H2), with minimal displacement along other axes. Consequently, a single static decision plane separating leftward and rightward choices would generate different fractions of leftward and rightward choices in the two contexts, accounting for the observed behavioral biases.

This geometric account was supported by neurometric analyses of the lPFC population activity patterns. A linear classifier trained to discriminate rightward and leftward choices in both contexts showed systematic shifts in output (Fig. 3c; Δ*β*_0_ = 1.5±0.6, *P* = 3.8×10^−3^, non-parametric one-sided bootstrap test): the probability of a test trial being classified as a rightward choice was larger on context right (blue curve) than on context left (green curve), paralleling the observed context-dependent behavioral bias (Fig. 2a). The distance of the lPFC activity patterns to the classifier’s discriminant hyperplane quantified the magnitude of this shift in the full-dimensional activity space (Fig. 3d; Δ*β*_0_ = 1.1 ± 0.3, *P <* 10^−3^, non-parametric one-sided bootstrap test), indicating tangible displacements that matched the psychometric shifts (Fig. 2a).

To further understand the representational geometry and investigate its computational consequences, we asked how well a linear decoder trained on one context generalized to the other. Generalization between contexts indicates disentangled geometries and is commonly quantified using the Cross-Condition Generalization Performance (CCGP).^35–37, 39, 40^ Standard CCGP asks whether a classifier trained on one subset of conditions generalizes to others. Here, because manifolds were shifted, we extended CCGP into a shifted-CCGP (shCCGP) (Fig. 3e), where we computed the hyperplane that separated leftward and rightward choices in one context and determined how much better the same hyperplane did in the other context when it was properly shifted. The aim of this extension is not to discover the readout used by downstream circuits but to characterize the geometry. If manifolds were structured but only shifted, performance should improve by adjusting the bias term. Indeed, shCCGP’s choice prediction accuracy significantly exceeded standard CCGP (Fig. 3f) and had a performance close to a decoder specifically trained for the other context.

Both CCGP and shCCGP for choice rose above baseline ∼150 ms after motion onset, but the rate of rise was significantly higher for shCCGP compared to CCGP (*P* = 1.6 × 10^−2^, one-sided z-test). CCGP and shCCGP could also be applied for decoding context. Both remained significantly above baseline throughout the trial, but CCGP fell below shCCGP ∼150 ms after motion onset (*P* = 2.0 × 10^−2^, one-sided z-test), further supporting that a shift disentangled the manifolds. Equivalent conclusions were obtained with alternative metrics such as the parallelism score (Supplementary Fig. S5).

### The neural signature of context inference

Behavioral analyses showed that monkeys performed a model-based context inference (Fig. 2e-f) when switching between reward contexts. We tested whether this inference was reflected in lPFC population activity by training a linear classifier to decode reward context. In well-trained monkeys, decoding performance adapted rapidly following a context switch (Fig. 3g). Because context switches were not explicitly signaled, the first rewarded trial after a switch (trial index 0) was the first informative cue. The context-representing lPFC neurons immediately adapted their activity after this first unexpected reward such that on the next trial, the decoded reward context from the lPFC population decisively shifted away from the previous context and fully switched within 1-5 trials (Fig. 3g).

This adaptation generalized across choices after a single unexpected feedback. Both tested trials (same chosen target as trial index 0) and untested trials (the opposite choice) showed rapid alignment with the new context (Fig. 3h; tested *P* = 2.7×10^−3^, untested *P* = 6.0×10^−3^, one-sided t-test). This generalization mirrors the behavioral results (Fig. 2e) and indicates that lPFC representations serve as a neural substrate for context inference, doing so using a single unexpected feedback to update the monkey’s internal model of the task (Fig. 2f-h). Individual monkeys showed similar results (Supplementary Fig. S4).

### Simulated Recurrent Neural Networks reproduce the observed behavior and representational geometry

The representational geometry observed in lPFC emerges naturally in artificial Recurrent Neural Networks (RNNs) trained to perform the alternating-context direction discrimination task. Using backpropagation through time,^39, 51–53^ we trained small RNNs (*N* = 10) that received two external inputs: a one-dimensional stochastic input that mimicked fluctuations of motion energy in a random-dot stimulus, and a binary signal that represented the current reward context (Fig. 4a). The network state at time *t* (**h***_t_*) evolved based on the weighted sum of its state at the previous time step (**h***_t_*_−1_), the external inputs (**x***_t_*), and independent gaussian noise (*σ**ξ**_t_*), passed through a nonlinear activation function. Each trial consisted of 20 time steps, with the motion input at each moment drawn from a Gaussian process whose mean corresponded to motion coherence in the trial (ranging from −1 for strong leftward motion to +1 for strong rightward motion). The context input was either +1 or −1 for context right and left, respectively, and persisted throughout the trial. The network was trained to integrate the motion input and make a decision at the end of the trial that minimized a loss function. To mimic the asymmetric reward contingencies in our task, incorrect leftward choices within the Context Left trials incurred larger penalties than incorrect rightward choices, and vice versa for Context Right trials. With this simple cost function, trained RNNs reproduced the key behavioral signatures (Fig. 4b): they chose the higher-paying choice more frequently and faster in each context. RNN reaction times were defined as the time within a simulated trial when the decision variable crossed a given threshold. Increasing the loss asymmetries in the network systematically amplified these choice biases and RT differences (Supplementary Fig. S6), consistent with normative and mechanistic models of decision making.^11, 24, 25^

**Figure 4:**
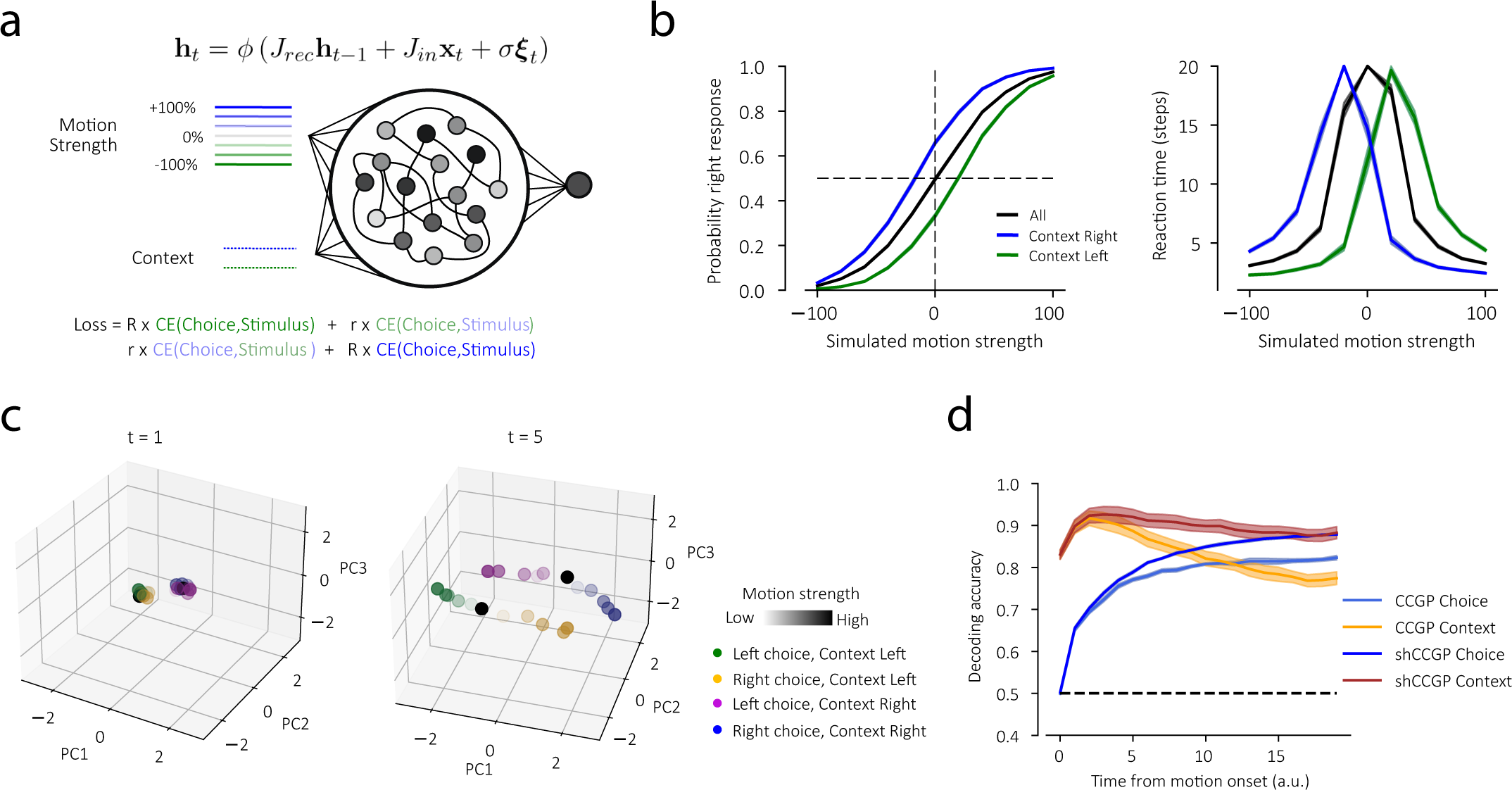
RNNs reproduce the behavioral and representational signatures of reward context switches. (**a**) Schematic of RNN architecture. Networks were trained to integrate a stochastic motion signal and a context signal to generate gain-maximizing choices. Context determined the asymmetric penalty for incorrect choices according to the loss function (r and R indicate asymmetric penalties, and CE is cross-entropy; see Methods). (**b**) RNN psychometric (left) and chronometric (right) functions for all trials (black), context left (green), and context right (blue). Networks chose the higher-reward option in each context more frequently and faster. Error bars show s.e.m. across 50 trained networks. (**c**) PCA projection of an example RNN population activity early (left) and late (right) in the trial. Late activity forms two shifted manifolds corresponding to the left and right contexts. (**d**) Comparison of standard CCGP and shifted-CCGP for direction choices (blue) and context (orange) across time within a trial. Error bars show s.e.m. across 50 networks. Results in b are shown for {*R* = 2*, r* = 1} and in c-d for {*R* = 4*, r* = 1}. See Supplementary Figs. S6, S7, and S8 for changes of RNN behavior and representational geometry with the magnitude of reward asymmetry.

We projected the RNN population activity onto its top three principal components (Fig. 4c). Early in the trial, different motion strengths were indistinguishable, and the representations collapsed on two clusters representing the two contexts. As motion input accumulated, activity patterns associated with different motion strengths unfolded along curvilinear manifolds within each context. Importantly, the two manifolds were shifted relative to each other along the decision variable axis, paralleling the geometry found in the neural data (cf. Fig. 3). The magnitude of this shift scaled with loss asymmetry, supporting a direct computational role for the representational displacement (Supplementary Fig. S7).

To further probe the geometry of representations in the full-dimensional state space of RNNs, we used cross-validated classifiers as we did for the neural responses (Fig. 4d). Linear classifiers trained on RNN population activity decoded choices with growing accuracy over time. The contexts were accurately decodable throughout the trial. Also, cross-condition generalization analyses yielded analogous results. The choice and context CCGP were significantly above chance, but the shifted-CCGP (shCCGP) exceeded the standard CCGP for both variables. Further, stronger reward asymmetries produced stronger shifts between the two manifolds as quantified by the shCCGP analysis (Supplementary Fig. S8).

These findings were robust across architectures, including GRUs, LSTMs, RNNs with different numbers of hidden units, and RNNs constrained by Dale’s law to have distinct excitatory and inhibitory units (Supplementary Fig. S9). Importantly, even in RNNs where the context signal was provided directly to the output unit of the networks (decision plane), they still exhibited shifted manifolds (Supplementary Fig. S10). For these modified RNNs, the behavioral bias was jointly implemented as a combination of a smaller shift in the representational plane and a shift in the decision plane. Overall, these results suggest that the representations observed in lPFC during the alternating-context emerge naturally from recurrent computations optimized for hierarchical decision-making under changing reward contexts.

### Behavioral and neuronal generalization throughout learning

Our experiments offered a unique opportunity to track how representations evolved as monkeys learned the task structure and inferred reward context switches. If lPFC representations underlie behavioral adjustments to reward bias, then neural context signals should evolve alongside behavior during training. Previous sections focused on the final third of recording sessions, when monkeys were fully trained. Here, we examine earlier sessions, when monkeys learned the alternating reward contexts. Before exposure to switching reward contexts, both monkeys were expert at the standard direction-discrimination task with symmetric rewards for the two choices. Electrophysiological recordings began with the introduction of asymmetric reward contexts, allowing us to track learning-related changes in both behavior and neural representations. Across training, behavioral performance and lPFC representations changed systematically and in tandem (Figs. 2c and 3g).

For each training session, we quantified the behavioral and neuronal *switch thresholds*. Behavioral threshold was defined as the number of trials the monkey required to switch its dominant choice (crossing 0.5 in Figs. 2c). Neural thresholds were defined analogously, as the number of trials required for neurally-inferred context to match the current context (crossing 0.5 in Figs. 3g). Behavioral thresholds decreased progressively across training (green line, Fig. 5a), indicating faster behavioral adaptation following context switches (Early vs. Late, *P* = 8.9 × 10^−3^, one-sided t-test). In the final sessions, adaptation occurred within only a few trials (∼3 trials for Monkey 1 and ∼5 trials for Monkeys 2). Neural thresholds showed a parallel reduction (blue line, Fig. 5a), revealing a close correspondence between behavioral and neural adaptation (Early vs. Late, *P* = 2.7 × 10^−5^, one-sided t-test). Consistent with this relationship, behavioral and neuronal thresholds were significantly correlated across sessions (Fig. 5b; Pearson *ρ* = 0.69, *P* = 1.1 × 10^−4^, two-sided t-test).

**Figure 5:**
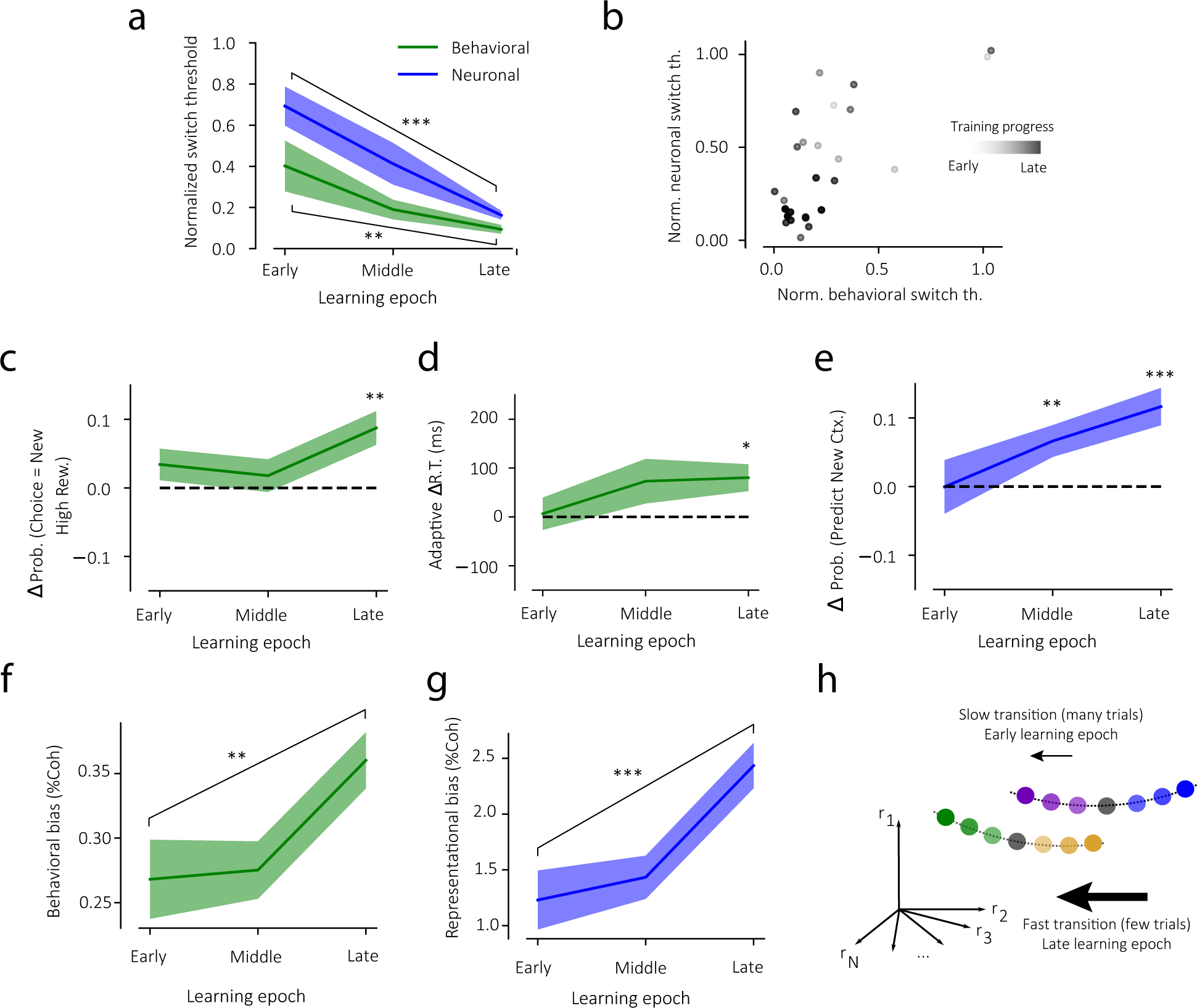
Monkeys learn to generalize more rapidly, supported by the geometry of lPFC representations. **(a)** Behavioral (green) and neural (blue) switch thresholds throughout learning. Behavioral thresholds indicate the number of trials required for the probability of choosing the new high-reward target to exceed 0.5 after a context change; neural threshold indicate the number of trials required for lPFC context signals to switch. The two thresholds decreased in tandem with training. Thresholds were normalized and pooled across monkeys. Early, middle, and late indicate the first, middle, and final thirds of training sessions. See Supplementary Fig. S11 for individual monkeys. (**b**) Correlation between behavioral and neuronal thresholds across training sessions. Gray shading denotes training progress. (**c**) Improvement in context inference from a single reward feedback across training, measured as the change in the probability of choosing the new high-reward target on trial index 1 relative to pre-switch trials. (**d**) Reaction-time adjustments on trial index 1 after a context switch, showing faster adaptation with training. (**e**) Change in neural decoding probability of context on trial index 1 after a switch, revealing faster representational transitions in later sessions. (**f**) Change in behavioral bias on trial index 1 after a switch, expressed in units of motion strength (*β*_2_*/β*_1_, Eq. 8). (**g**) Corresponding bias derived from lPFC neural representations. (**h**) Schematic summary: early in training, lPFC population activity required multiple trials to switch context representation; late in training, this transition occurred faster, often after a single feedback. Shaded regions in a, c-e indicate s.e.m. across sessions. Shaded regions in f-g indicate standard error of the logistic fit (see Methods). ∗ = *P <* 0.05, ∗∗ = *P <* 0.01, ∗ ∗ ∗ = *P <* 0.001

Direct analyses of choice bias and response time further showed that the ability to infer context from a single unexpected reward—a hallmark of model-based generalization (Fig. 2e,f)—emerged gradually during training. Early in training, monkeys required several rewarded trials to adjust reaction times following a context switch, whereas in later sessions, they demonstrated significant adjustments of reaction times for the untested target on trial index 1 (Fig. 5d). Similarly, the probability of choosing the new high-reward target on trial index 1 increased across training (Fig. 5c). The rate of post-switch bias adjustment increased significantly across training (*β*_3_ *>* 0, Eq. 7, *P <* 10^−8^, likelihood ratio test), leading to faster shifts in the psychometric function following context switches (Fig. 5f; early vs. late change in bias, (9.2 ± 3.7) × 10^−2^,*P* = 7.1 × 10^−3^, one-sided z-test).

Prearcuate population activity showed a corresponding adaptation. Early in training, neural representations required several trials to align with the new context, whereas in late sessions they switched almost immediately after a single unexpected reward (Fig. 5e; see Supplementary Fig. S12 for changes of individual unit response dynamics). Similarly, neural choice bias adjusted faster across training (*β*_3_ *>* 0, Eq. 7, *P <* 10^−8^, likelihood ratio test), yielding larger biases on trial index 1 in late training (Fig. 5g; change of bias, 1.2 ± 0.3,*P* = 1.4 × 10^−4^, one-sided z-test).

Although early in training both behavioral and neural responses required many trials to fully adjust after a context change, the final magnitude of the bias reached within each block was comparable throughout learning (Supplementary Fig.S13). Thus, learning primarily altered the speed of adaptation rather than its asymptotic level. Over training, monkeys progressively learned to infer context switches from a single feedback, and lPFC neural representations evolved in lockstep with these behavioral changes (Fig.5h).

## Discussion

Adaptive behavior in dynamic environments requires integrating sensory information with contextual knowledge about task structure. In the alternating-context direction-discrimination task, monkeys learned to combine momentary evidence with reward context to maximize payoff. They biased their choices toward the more rewarded option in each context and, with experience, learned to adjust that bias after a single feedback—an instance of task inference. Neural recordings from lateral prefrontal cortex revealed that this inference was accompanied by systematic shifts in the geometry of population activity: stimulus information was represented on two curvilinear manifolds corresponding to the two contexts, displaced relative to each other along the decision-variable axis. This representational shift provided a simple geometric mechanism for implementing context-dependent biases through a fixed linear readout. Moreover, the emergence of faster inference at the behavioral level was mirrored by faster transitions between the two manifolds in lPFC population activity, linking learning of task structure to the evolution of neural representations.

Importantly, our goal was not to teach monkeys how to make decisions under asymmetric rewards—a skill they likely possessed through out-of-task experiences—but to study how they learned the structure of a novel environment in which reward contingencies changed without explicit cues. Learning in this setting required forming and updating an internal model of context transitions and using single outcomes to infer latent states. The behavioral and neuronal signatures of this inference—context generalization, rapid bias adjustment, and representational switching—reveal how prefrontal circuits support hierarchical forms of learning that extend beyond stimulus–response mapping.

The chronic stability of our recordings allowed us to track neural activity within the same cortical micro-columns across the entire training period. While our analyses did not rely on continuous tracking of individual neurons, stable multi-day recordings ensured that changes in representational geometry reflected adaptation within a population rather than recruitment of new neural sub-populations.^54, 55^ This stability strengthens the conclusion that learning to infer context was accompanied by true changes in the representational format of prefrontal neurons.

At the population level, lPFC activity evolved along low-dimensional manifolds whose relative displacement encoded reward context. These shifts paralleled the behavioral biases observed across contexts and provided a natural substrate for flexible inference. Shifting manifolds along the decision-variable axis allows downstream readouts to adjust their output without requiring synaptic changes or explicit context signals. This transformation differs from previously reported geometric reorganizations during rule changes, in which manifolds either translate orthogonally to the decision axis^35, 45, 51, 56^ or rotate in state space, requiring context-dependent readouts.^41^ In contrast, context-dependent reward asymmetries produce mainly a shift along the decision-variable axis. By shifting activity along this axis, prefrontal populations can recalibrate policy while preserving the same evidence-to-choice readout, providing a mechanistic bridge between rapid, outcome-driven inference and stable downstream decoding. These results outline a compact geometric account of how prefrontal circuits learn and deploy latent context to control decisions—updating behavior after minimal feedback without rebuilding the decision machinery. Further, adding shifts to rotations and orthogonal translations forms a compact vocabulary for understanding how neural populations reorganize to support adaptive behavior.

Recurrent neural network models trained on the same task reproduced the key representational features observed in lPFC. Without explicit architectural constraints, these networks developed parallel, context-dependent manifolds whose displacement scaled with reward asymmetry. This suggests that the observed geometry is a natural computational solution for distributed systems learning to optimize performance under changing reward contingencies. Similar representational shifts are likely to arise in other situations that require changes of decision bias, such as those induced by priors^3, 11, 12, 17^ or by heuristics that modulate confidence and decision thresholds.^13, 28^

Our findings position the lateral prefrontal cortex within a hierarchical decision-making system that not only integrates sensory evidence but also dynamically infers and applies contextual knowledge. In this framework, context representations act as a control layer that configures the mapping from evidence to action^2^—essentially gating decision policy rather than implementing it. Indeed, recent work has shown that thalamocortical pathways can reconfigure prefrontal circuits during flexible switches in task structure, by promoting rapid realignment of cortical representations.^57, 58^ Shifting low-dimensional population manifolds along decision-variable axes enables circuits to adapt policy without re-learning stimulus–response mappings, thereby offering both flexibility and efficiency. More broadly, the geometric operations observed in frontoparietal networks may reflect a general mechanism by which internal models of latent context are instantiated and applied.

## Methods

We recorded neuronal populations in the prearcuate gyrus (area 8Ar) of two adult macaque monkeys (Macaca mulatta; male, aged 7-9 years) performing a variant of the motion direction-discrimination task with random dots. All procedures conformed to the National Institutes of Health *Guide for the Care and Use of Laboratory Animals* and were approved by the Institutional Animal Care and Use Committee at New York University.

### The alternating-context direction-discrimination task

Monkeys were seated in a semi-dark room in front of a CRT monitor (frame rate, 75 Hz) with heads stabilized using surgically implanted titanium headposts. A custom Matlab (Mathworks Inc., Massachusetts, USA) program controlled stimulus presentation and task flow using Psychophysics Toolbox.^59^ We monitored the monkey’s gaze at 1 kHz using a high-speed infrared camera (Eyelink, SR-Research, Canada).

Fig. 1 illustrates task design. Each trial began when the monkey fixated a central fixation point (FP, 0.3^◦^ diameter). After a variable delay (150-450 ms; truncated exponential distribution), two red targets appeared: one contralateral to the recorded hemisphere, aligned with the response fields (RFs) of most recorded neurons (eccentricity, 10.5^◦^ for monkey 1; 15^◦^ for monkey 2), and the other symmetrically placed on the opposite side of the screen. RF locations were determined using a memory-guided saccade task performed in separate blocks. Following an additional delay (350-550 ms; truncated exponential distribution), a random-dot motion stimulus^60, 61^ appeared within a circular aperture centered on the FP (diameter, 5^◦^). The stimulus comprised three interleaved sets of dots displayed on alternating frames. Each set was shown for one video frame (13.3 ms) and updated every three frames (Δt = 40 ms). A subset of dots moved coherently at 5 ^◦^/s in the same direction, while the rest were replotted in random locations (dot density, 16.7 dots/deg^2^/s). Motion direction (left or right) and motion strength remained constant within a trial and varied randomly across trials. Motion strengths (0%, 1.6%, 3.2%, 6.4%, 12.8%, 25.6%, 51.2%) were chosen to span the full performance range, from chance to perfect accuracy. Monkeys could report the perceived motion direction any time after motion onset by making a saccade to one of the targets. Correct choices yielded liquid reward. For 0% motion strength, the monkey was rewarded randomly on half of the completed trials irrespective of the choice. Trials were aborted if the gaze deviated by more than 2^◦^ from the FP before stimulus onset or if gaze shifts occurred outside of a saccade to a target. Distinct auditory feedback signaled correct, incorrect, or fixation-break outcomes.

After mastering the basic direction-discrimination task, we implanted recording electrodes and introduced the alternating reward contexts. In context R, correct responses to the right target yielded larger reward than to the the left target; in context L, the reward asymmetry was reversed. Reward contexts persisted for tens to hundreds of trials before switching stochastically and without explicit cues. Reward ratios (higher-reward / lower-reward) were 1.22 and 1.5 for monkeys 1 and 2, respectively. Initial sessions featured slightly higher ratios to attract the monkey’s attention to the task change. Reward ratios were then reduced to the final values. The data analyzed in the present study include all sessions with reward imbalance, from early training to the final, well-trained stage: 12 sessions from monkey 1 (17,957 trials) and 30 sessions from monkeys 2 (43,556 trials). For the last third of the sessions—used for “expert” analyses—context switches occurred every 135 trials on average for monkey 1, and every 265 trials for monkey 2.

### Neural recordings

Before introducing the asymmetric reward contexts, we implanted 96-channel microelectrode arrays (electrode length, 1 mm; inter-electrode spacing, 0.4 mm; ∼ 0.5*M* Ω impedance; Blackrock Microsystems, UT) in the prearcuate gyrus (Fig. 3). While monkeys performed the task, neural spike waveforms were recorded (sampling rate, 30 kHz) and aligned to task events for subsequent analyses. We performed offline sorting for a subset of sessions (Plexon Inc., Dallas, TX), but observed qualitatively similar results for the sorted and unsorted spikes in our analyses, consistent with previous reports demonstrating the sufficiency and robustness of threshold-crossing activity for population analyses.^62^ Consequently, we conducted all main analyses using unsorted multi-unit activity to reduce unnecessary preprocessing and enhance robustness of analyses. Throughout the paper, we use ‘units’ to refer to multi-unit activity recorded from individual electrodes. All units were retained in our analyses irrespective of selectivity. PCA results for spike-sorted expert sessions of individual monkeys are shown in Supplementary Figs. S2 and S3.

### Analysis of Behavior

We used a logistic function to explain the relationship between the probability of a rightward choice and motion strength (% Coherence) (Fig. 2a):

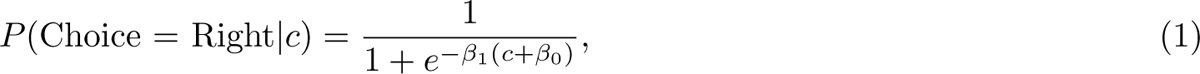

where *c* is the signed motion strength (positive for rightward, negative for leftward motion), and *β*_1_ and *β*_0_ are the slope and bias parameters. We fit three separate psychometric functions for right-context trials (blue), left-context trials (green), and all trials combined (black).

For chronometric analyses (Fig. 2b), we computed the mean reaction time (RT) for each motion strength and context of each session. We used a hyperbolic tangent function to explain changes of mean RT with stimulus strength:^46, 63^

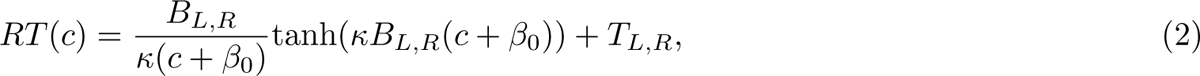

where *c* is signed motion strength, *κ* is the sensitivity parameter (related to *β*_1_ in Eq. 1), *β*_0_ capture bias (related to *β*_0_ in Eq. 1), and *T_L,R_*and *B_L,R_* are the non-decision time and evidence accumulation bounds, respectively. To capture potential reaction time asymmetries, we allowed different *T* and *B* for the two choices.

Figure S1a,b shows the mean and s.e.m. of fits across the final third of sessions, when monkeys performed at peak proficiency (monkey 1, *n* = 4*/*12 sessions; monkey 2, *n* = 10/30 sessions). Figure 2a,b shows behavior pooled across monkeys. Combined errorbars were calculated by pooling uncertainty: 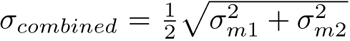, where *σ_m_*_1_ and *σ_m_*_2_ corresponded to the s.e.m. for each monkey. For each session, we computed context-dependent change of bias, Δ*β*_0_, as the difference in fitted *β*_0_ between right and left contexts. The reported values correspond to the mean ± s.e.m. across sessions from both monkeys. Statistical significance was assessed with two-sided paired t-tests.

### Denoising of population activity and PCA

To visualize the geometry of neural responses across motion strengths and reward contexts, we first denoised single-trial firing rates using an encoding model and then projected the denoised population activity onto its top 3 principal components (see [39]). The model was optimized to predict the firing rates:

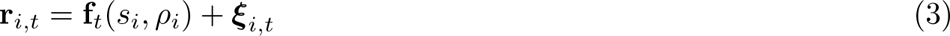

where **r***_i,t_* is the population firing rate vector on trial *i* at time *t*, *s_i_* is the signed log-transformed motion strength (sign(*c_i_*) × log(|*c_i_*|)), *ρ_i_* is reward context (−1 and +1 for context L and R), and ***ξ****_i,t_* is i.i.d. Gaussian noise (independent across neurons and time-steps). For the encoding model, **f***_t_*, we used a feedforward artificial neural network with an input layer, two hidden layers with 100 ReLU units per layer, and a linear output layer. This model was chosen based on cross-validation performance of networks with zero to four hidden layers (the zero-layer network is equivalent to linear regression). We trained the model using z-scored firing rates of neurons in 200 ms windows, minimizing the mean squared error between the normalized firing rates of recorded neurons and model predictions (stochastic gradient descent implemented with MLPRegressor of sklearn library; learning rate, *lr* = 10^−3^; L2 regularization *λ* = 10^−3^).

We then projected the denoised population activity onto the top three principal components. Firing rates for each motion strength and reward context were computed using a sliding 200 ms window. The top three PCs were learned from a data matrix that contained firing rates from all motion strengths and contexts throughout the stimulus viewing epoch. To ensure that the visualization of representational geometries are not contaminated by the transient neural activity around the response saccade, time windows overlapping the last 50 ms before saccade onset were excluded from each trial. The sliding window advanced from motion onset until less than 25% of trials contributed to the calculation of averaged firing rates. Figure 3b shows PCA projections from a representative session at two time windows: an early interval (*t* = [0, 200] ms) after motion onset, when the integration of evidence was not yet represented by prearcuate neurons, and a later interval (*t* = [400, 600] ms) when evidence integration was underway but before saccade initiation. Supplementary Figs. S2,S3 show additional example sessions from each monkey.

We used a similar analysis pipeline to visualize the geometry of RNN activity (no denoising step was necessary for RNNs). Fig. 4c shows projections of the activity of a representative network at early and intermediate times, similar to Fig. 3a (*t* = 1 and *t* = 5). See Supplementary Fig. S7 for additional networks, time points, and reward ratios.

### Cross-Condition Generalization Performance (CCGP) and shifted-CCGP

To determine the representational structure of prearcuate population activity, we trained classifiers to decode motion-direction choice, reward context, and their XOR conjunction from neural population activity, and then used the classifier performance and generalization behavior to quantify representational geometry. Firing rates were computed in sliding 200 ms windows up to 50 ms before saccade onset. Analyses were conducted on individual sessions or pseudo-populations constructed across sessions. Here, we focus on pseudo-populations as they provide a simpler analysis pipeline, though single-session analyses yielded qualitatively similar results. We constructed pseudo-populations by sampling correct trials with similar signed motion strengths and reward context across sessions (100 trials per condition). To eliminate train-test overlap, trials from each session were first partitioned into distinct train (80%) and test (20%) sets, and pseudo-populations were assembled independently within each set. Each session’s firing rates were z-scored before concatenation across sessions.

At each time bin after motion onset, we fit linear classifiers (logistic regression with L2 regularization, *λ* = 100) to decode motion-direction choice, reward context, or their XOR. The XOR classifier discriminated trial group {right choice/context R, left choice/context L} from {right choice/context L, left choice/context R} and its performance was diagnostic of representational dimensionality.^35, 43^ For each analysis, we repeated pseudopopulation construction and classifier training/testing 10 times, and calculated the mean and standard deviation of classifier performance across these repetitions. Statistical significance was assessed using null distributions generated by shuffling trial labels for motion strength and context within sessions, repeating pseudopopulation construction and decoding 10 times per shuffle. This process was repeated 1000 times to make the null distribution for decoder performance. Shaded areas in Supplementary Figure S4c correspond to the 2.5th—97.5th percentiles of these null distributions.

We used these classifiers in five analyses. First, we used the cross-validated choice prediction accuracy to quantify the “neurometric” curves for each reward context (Figs. 3c and S4a). The classifiers were trained and tested using the mean firing rates of neurons throughout stimulus viewing, excluding the final 50 ms prior to saccade onset. To quantify changes of neurometric bias across reward contexts, we fit the logistic function in Eq. 1 to cross-validated accuracies for different motion strengths and computed Δ*β*_0_ = *β*_0*,R*_ − *β*_0*,L*_. Motion strength was signed log-transformed before fitting Eq. 2 to improve our power for identifying small shifts. A bootstrap test was used to quantify significance of Δ*β*_0*,i*_ *>* 0 (1000 iterations).

Our second class of analyses focused on the distance of the population activity patterns from the choice classifier’s discriminant hyperplane (Figs. 3d and S4b): *d* = ***ω****^T^* **x** + *ω*_0_. This distance quantifies the divergence of neural responses as a function of time, choice, and stimulus strength, providing a neural proxy for the evolving decision variable.^13, 50, 64, 65^ We quantified the change of distance with reward context by regressing the signed distance against signed log-transformed motion strengths, and calculated intercept changes. Statistical significance was assessed using a bootstrap test (1000 iterations).

The third and fourth classes of analyses directly targeted the geometry of representations. CCGP characterizes representational disentanglement: whether a classifier trained in one subset of conditions generalizes to another.^35, 39^ For choice CCGP, classifiers trained to discriminate right and left choices in one reward context were tested in the other. For context CCGP, classifiers trained to discriminate contexts for one choice were tested on the other choice. To isolate geometric shifts between manifolds, we developed shifted-CCGP (shCCGP). If the two contexts differ mainly by a translation of representational manifolds, then we should be able to increase CCGP by adjusting the classifier’s bias parameter *ω*_0_. We therefore scanned decoding performance for different values of *ω*_0_ and picked the bias that maximized generalization performance (Figs. 3e-f and S4d; see also Supplementary Fig. S5). shCCGP was computed for both the choice and context decoders. Error bars in the figures correspond to standard deviation of CCGP and shCCGP across 10 repetitions of pseudo-population reconstructions.

Chance performance for CCGP and shCCGP were generated following the method described in [35, 39]. Briefly, we rotated pseudopopulation activity patterns to random locations in the neural state space by condition-wise shuffling of neuron identities. This rotation was consistent for all the trials of the same condition (a given choice and context), but varied across conditions, destroying any representational structure. Then, CCGP and shCCGP were recomputed as explained above. The shaded areas reported in Figures 3f and S4d correspond to the 2.5th-97.5th percentiles of the null CCGP and shCCGP distributions obtained by repeating this process 1000 times.

Our fifth analysis calculates parallelism scores (Supplementary Fig. S5d) following [35]. We computed the cosine similarity between (1) reward context decoders trained separately on left-choice vs. only right-choice trials, and (2) choice decoders trained separately context R vs. context L trials. High parallelism scores indicate aligned representational axes across conditions. Null distributions were generated using the same random-rotation procedure described above.

### Recurrent neural network

We trained recurrent neural networks (RNNs) to perform the alternating-context direction-discrimination task. Each RNN consisted of *N* = 10 units whose activity at time *t* (**h***_t_*) was determined by the following equation:

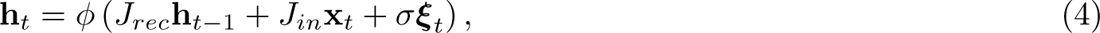

where *ϕ*(.) is the tanh non-linearity, ***ξ****_t_* is i.i.d. unit-variance Gaussian noise on each time step, *σ* = 1 is the noise amplitude, and *J_rec_* and *J_in_* are recurrent and input weight matrices, respectively. The input **x***_t_* included momentary sensory evidence (fluctuations of motion) and reward context. The momentary evidence on trial *i* was simulated as a Gaussian process with mean *α_i_* and standard deviation 1. Each trial consisted of up to 20 time steps (∼75 ms each). Values of *α_i_* determined stimulus strength and were uniformly sampled from [−100, −80, −60, −40, −20, 0, 20, 40, 60, 80, 100], with negative and positive values corresponding to leftward and rightward motion, respectively. Values near ±1 generated easy stimuli, whereas values near 0 produced difficult ones. The context input was +1 (context R) or −1 (context L), held constant throughout the trial, with additive Gaussian noise with standard deviation 1 to mimic potential memory fluctuations.

A linear readout unit produced an output *O_t_* approximating the integrated evidence, with a context-dependent bias to maximize expected reward. We defined reaction time on each trial as the first time step when |*O_t_*| reached a criterion level (0.7 for asymmetric reward contexts and 0.9 for symmetric contexts).

RNNs were trained to minimize a context-dependent cross-entropy loss function:

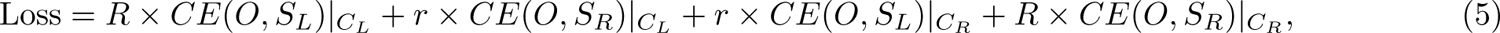

where *CE* denotes cross-entropy, *C_L_* and *C_R_*are leftward and rightward contexts, and *S_L_* and *S_R_*are leftward and rightward motion directions. Reward asymmetry was controlled by the pair [*R, r*], which was set to [1, 1], [2, 1], or [4, 1] for symmetric, asymmetric, and strongly asymmetric environments. RNNs were trained using backpropagation through time (ADAM optimizer, learning rate 0.01) for 200 epochs with minibatches of 1000 trials and L2 regularization *λ* = 10^−5^. After training, each network was evaluated on a separate set of 2200 trials (200 trials per motion strength).

RNN psychometric and chronometric curves (Figs. 4b and S6b,e) were computed analogously to the behavioral analyses (Fig. 2a,b). For each signed motion strength, we computed the probability of the RNN choosing the rightward target (psychometric function) and the mean number of time steps for the decision variable to reach the decision bound (chronometric function). The reported means and s.e.m. were calculated across 50 independently trained networks.

RNN decoding, CCGP, and shCCGP analyses followed the same procedures as those used for neural activity. For each simulated session and time step, we first subsampled trials without replacement such that all four experimental conditions (leftward/rightward choice × context L/context R) were equally populated and thus direction choice and reward context became decorrelated. We split trials in each condition into 80% training and 20% testing sets. We trained three independent classifiers (choice, context, XOR) using logistic regression and evaluated them using five-fold cross-validation. For CCGP and shCCGP, we tested generalization performance of the choice or reward context classifiers for each time step and network as explained for the neural data. shCCGP adjusted the classifier bias *ω*_0_ to maximize cross-condition performance. The reported means and s.e.m. were calculated across 50 independently trained networks (Fig. 4d and Supplementary Fig. S8).

Supplementary Figure S9 demonstrates that the task solution and representational geometry were consistent across RNN architectures, including RNNs with *N* = 3 and *N* = 50 units (panels a,b), LSTM^66^ (panel c), and GRU^67^ (panel d) recurrent networks, and RNNs constrained by Dale’s law (panel e). For Dale-constrained RNNs, we added an additional penalty term to Eq. 5 to enforce two unit subpopulations: excitatory units with positive outgoing weights and inhibitory units with negative outgoing weights, with a target ratio of 4:1.

### Learning task model and inferring context switches

We tracked how the monkeys’ behavior and neural responses evolved from the first session in which asymmetric reward contexts were introduced. Prior to this, monkeys were trained only on the basic direction-discrimination task with symmetric rewards. We explored whether, across session, monkeys learned an internal model of reward-context transitions and whether they used this model to adapt behavior immediately following a context change. We recorded 12 sessions in monkey 1 and 30 sessions in monkey 2 throughout learning. For each monkey, we divided sessions into three learning epochs: early, mid, and late (monkey 1: sessions 1-4, 5-8, 9-12; monkey 2: sessions 1-10, 11-20, 21-30). The analyses reported in Figures 2 and 3 focus on the late epoch. Figure 5 shows changes of behavior and neural representations across epochs.

### Context inference following a switch

A change in reward context alters the expected rewards for both choice targets. An agent that has learned the task model should update expectations for both targets immediately after the first inconsistent reward, not only for the target chosen on that trial (“tested” target). For example, if the monkey has been in a left-context block (large rewards for left choices) but receives a large reward on a rightward choice, it should immediately infer that the reward context has switched and left choices will now yield small rewards, even though it has not yet chosen leftward (“untested” target) in the new context.

To test this, we first focused on the last third of sessions, when both monkeys had mastered the task. We defined the subjective context switch as the first correct trial following an objective context change, which yielded a reward inconsistent with the previous context (Fig. 2d). This trial was assigned index 0, and the chosen target on this trial was labeled tested. We then examined whether the probability of choosing the untested target and the associated RTs changed on subsequent trials in the direction predicted by the task model.

Reaction times were especially informative, as they provided a graded measure more sensitive than binary choices. For each switch, we computed the change in RT on trial index 1 relative to a pre-switch baseline (Adaptive ΔRT; Fig. 2f). The pre-switch baseline was obtained by fitting the chronometric function of Eq. 2 to the last 30 trials of the preceding block and calculating the expected mean RT for trials with the same motion strength and choice as trial index 1. Results reported in Fig. 2f correspond to the mean and s.e.m. across four sessions for Monkey 1 and five pairs of sessions for Monkey 2. Consecutive session pair were combined for monkey 2 to match the number of switches experienced by monkey 1 in each session. Statistical significance of RT adaptations for tested and untested choices on trial index 1 was evaluated using a one-sided t-test (Fig. 2f).

To quantify how bias evolved after a switch, we fit a sigmoidal function to the monkey’s choices across post-switch trials 1-80:

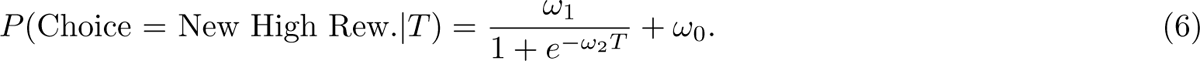

where *T* is trial index and *ω_i_* are model parameters. For each session (Monkey 1) and pairs of sessions (Monkey 2), we fit an independent function to the mean *P* (Choice = New High Rew.|*T*) across context switches for each combination of tested/untested and high/low reward. We computed ΔProb.(Choice = New High Rew.) as the fit value at *T* = 1 minus the mean probability of making the same choice in the 20 pre-switch trials. Results in Fig. 2e correspond to the mean and s.e.m. across sessions and reward magnitudes. We tested whether the distribution of ΔProb.(Choice = New High Rew.) was significantly above zero using a one-sided t-test.

We next quantified how quickly the prearcuate population activity reflected the new context. For each session, we trained a linear classifier to decode context from stimulus-aligned neural activity (monkey 1, 0-200 ms; monkey 2, 0-300 ms), using all trials except those around the context switch (−20 ≤ *T* ≤ 80). We then projected population activity patterns of the peri-switch withheld trials onto the classifier axis to compute trial-by-trial changes of predicted context. We fit Eq. 6 to these decoder outputs within trials 1-80 and defined ΔProb. (Predict New Context) analogously to the behavioral measure (Fig. 3h). Results for individual monkeys are shown in Supplementary Figures S1 and S4.

### Adaptive behavior across learning

We quantified how adaptation after a switch improved across learning. For each monkey, we fit the following logistic model to choices from trials −20 to +30 around context switches:

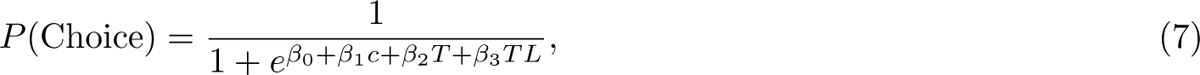

where *c* is signed motion coherence, *T* is trial index relative to the switch, and *L* is the cumulative number of context switches experienced across sessions—a proxy for learning progress. The interaction term *β*_3_*TL* captures whether adaptation became faster with experience. Separate fits were performed for *L* → *R* and *R* → *L* switches (monkey 1, 48 and 49; monkey 2, 62 and 59). Statistical significance of *β*_3_ was assessed using a likelihood ratio test against a model with *β*_3_ = 0.

To directly compare context learning across epochs (Fig. 5f), we also fit a reduced model without *β*_3_ to data from each learning epoch:

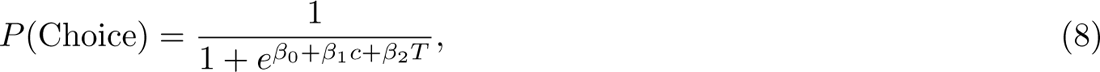

and plotted *β*_2_*/β*_1_—the rate of post-switch bias change in units of stimulus strength (% coherence). Statistical significance was assessed using a one-sided z-test against the early epoch.

To compare changes of adaptive ΔRT across training (Fig. 5d), we combined adaptive ΔRTs for high/low reward and tested/untested choices after flipping the sign of ΔRT for high-reward choices. We tested the significance of pooled values against zero using a one-sided Wilcoxon test.

To investigate changes in neural representations, we first trained a linear classifier to predict the monkey’s choice from population activity after motion onset (monkey 1, 0-400 ms; monkey 2, 0-600 ms). We excluded peri-switch trials for training and used the model to predict the monkey’s choice on the withheld trials. These neurally predicted choices reflected context-dependent biases (Fig. 3c-d). The full and reduced logistic models of Eqs. 7 and 8 were fit to these predicted choices as above. We used a likelihood ratio test to check whether the neural representation of bias adapted with learning (*H*_0_: *β*_3_ = 0, Eq. 7). We further used a z-test to check whether the neural bias changed significantly from the early to middle and late learning epochs (Fig. 5g).

To obtain the behavioral switch thresholds (Fig. 5a,b), for each session of monkey 1 and pairs of consecutive sessions of monkey 2, we computed Prob. (Choice = New High Rew.) by fitting the following equation to choices made on post-switch trials *T* = [1, 80]:

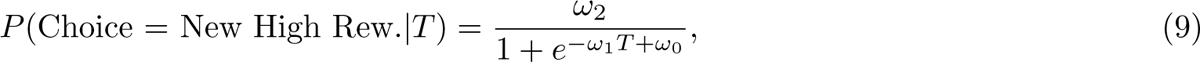

where *ω_i_* are model parameters. Fig. 2c shows the fit averaged across sessions for both monkeys. The pre-switch baseline was computed as mean of *P* (Choice = New High Rew.) across trials *T* = [−20, 0]. We defined the switch threshold as the trial number *T* ^∗^ in which the curve reached 0.5. Faster adaptations after a context switch will be associated with smaller *T* ^∗^ than slower adaptations. In Supplementary Fig. S1, we show the results for each monkey.

We performed an analogous analysis on neural representations to obtain neural switch thresholds. We first trained a linear classifier to predict the reward context from population activity after stimulus onset (monkey 1, 0-200 ms; monkey 2, 0-300 ms). We excluded peri-switch trials (*T* = [−20, 80]) for training and used the model to predict the context represented in lPFC on each trial around each context switch for −20 ≤ *T* ≤ 80. We fit Eq. 9 to these decoder outputs (Fig. 3g) and defined neural thresholds as the number of trials required to reach 0.5.

We plot behavioral and neuronal thresholds across learning epochs in Fig. 5a,b. Thresholds were normalized within each monkey by dividing by that monkey’s largest threshold throughout learning. We also computed the Pearson correlation between behavioral and neuronal thresholds and evaluated its significance with a two-sided, non-parametric permutation test (Fig. 5b). Further, we compared early vs. late thresholds using a one-sided t-test (Fig. 5a).

We also evaluated learning-dependent adaptation in individual units across sessions (Supplementary Fig. S12). For each neuron, we computed normalized firing rates in an early post-stimulus window (monkey 1, 0-200 ms; monkey 2, 0-300) from 50 trials before to 100 trials after each switch. The time course of single-unit adaptations was fit with a modified version of Eq. 9:

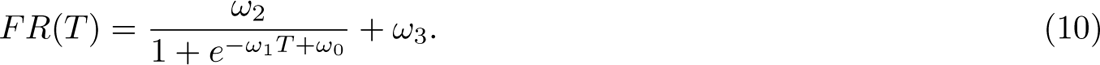

where *ω*_1_ quantifies the adaptation rate. Units with adaptation within *<*1 trial were excluded (monkey 1, 5.9%; monkey 2, 5.6%). Supplementary Figure S12 shows mean *ω*_1_ across neurons for early, mid, and late learning epochs.

## Acknowledgments

This work was supported by R01MH109180, R01MH127375, R01MH141929, the Pew Innovation Fund, R01NS094659, R01NS069679, F32NS096819, and U01NS099726. Additional support was provided by the Simons Foundation, NSF 1707398 (Neuronex), the Gatsby Charitable Foundation (GAT3708), and the Swartz Foundation. S.E. was supported by NYU’s Dean’s Thesis Fellowship Award.

## Author Contributions

R.K., S.F., and S.E. conceptualized and designed the study. S.E. and R.K. collected the data. R.N. performed the analyses and generated figures with guidance from R.K. and S.F. R.N. trained and analyzed the recurrent neural networks. R.N., S.F., and R.K. wrote the manuscript.

## Data availability

All datasets generated and analyzed in this study will be deposited in a public repository and made available upon publication.

## Code availability

The code is available at https://github.com/ramonnogueira/contextPFC.

## Supplementary Figures

**Figure S1:**
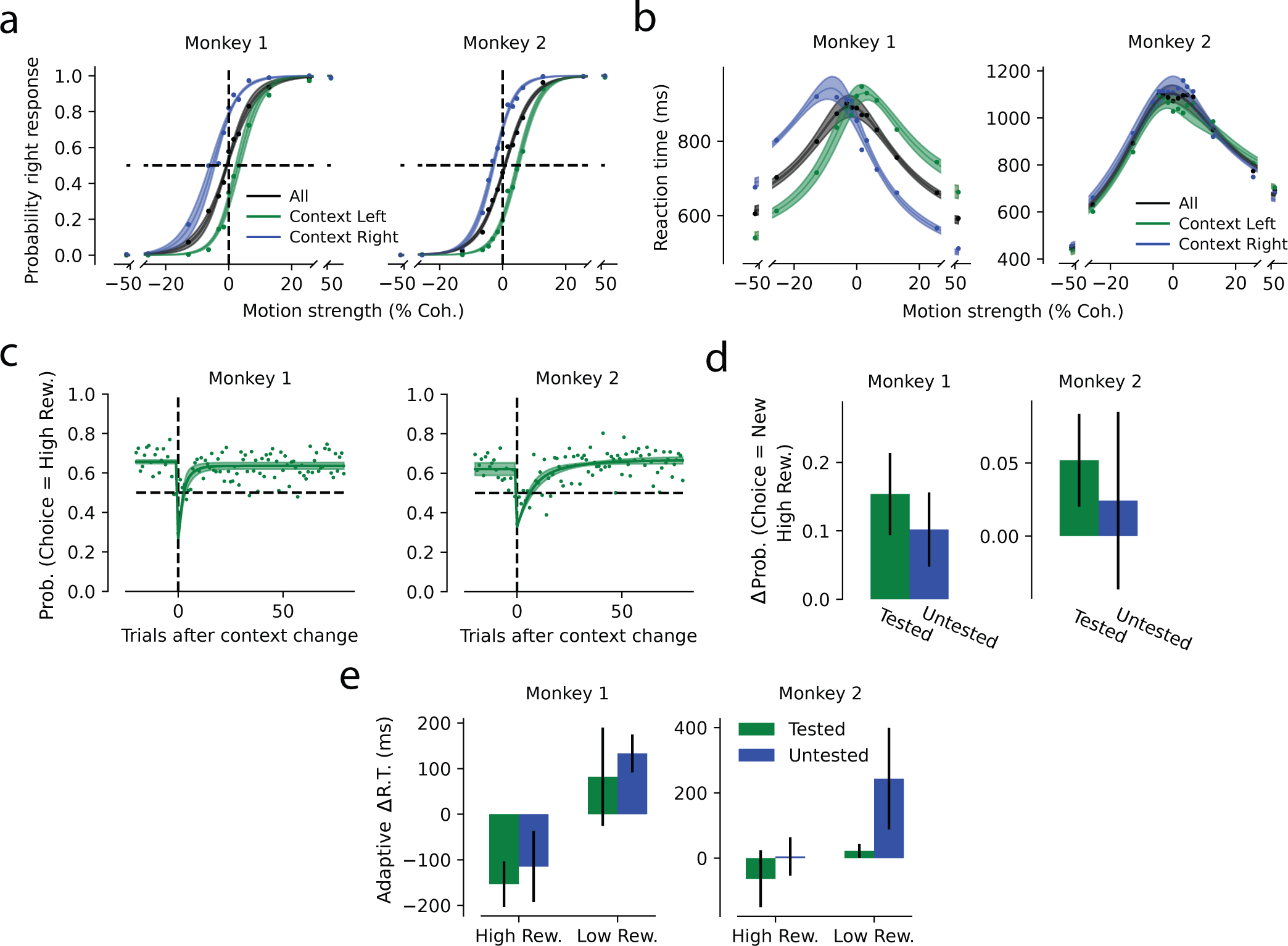
Behavioral signatures of reward bias and task inference in both monkeys.. Both monkeys biased their choices toward the higher-reward target in each context and adjusted reaction times following context switches. Panels correspond to those in Fig. 2 and are shown separately for each monkey.

**Figure S2:**
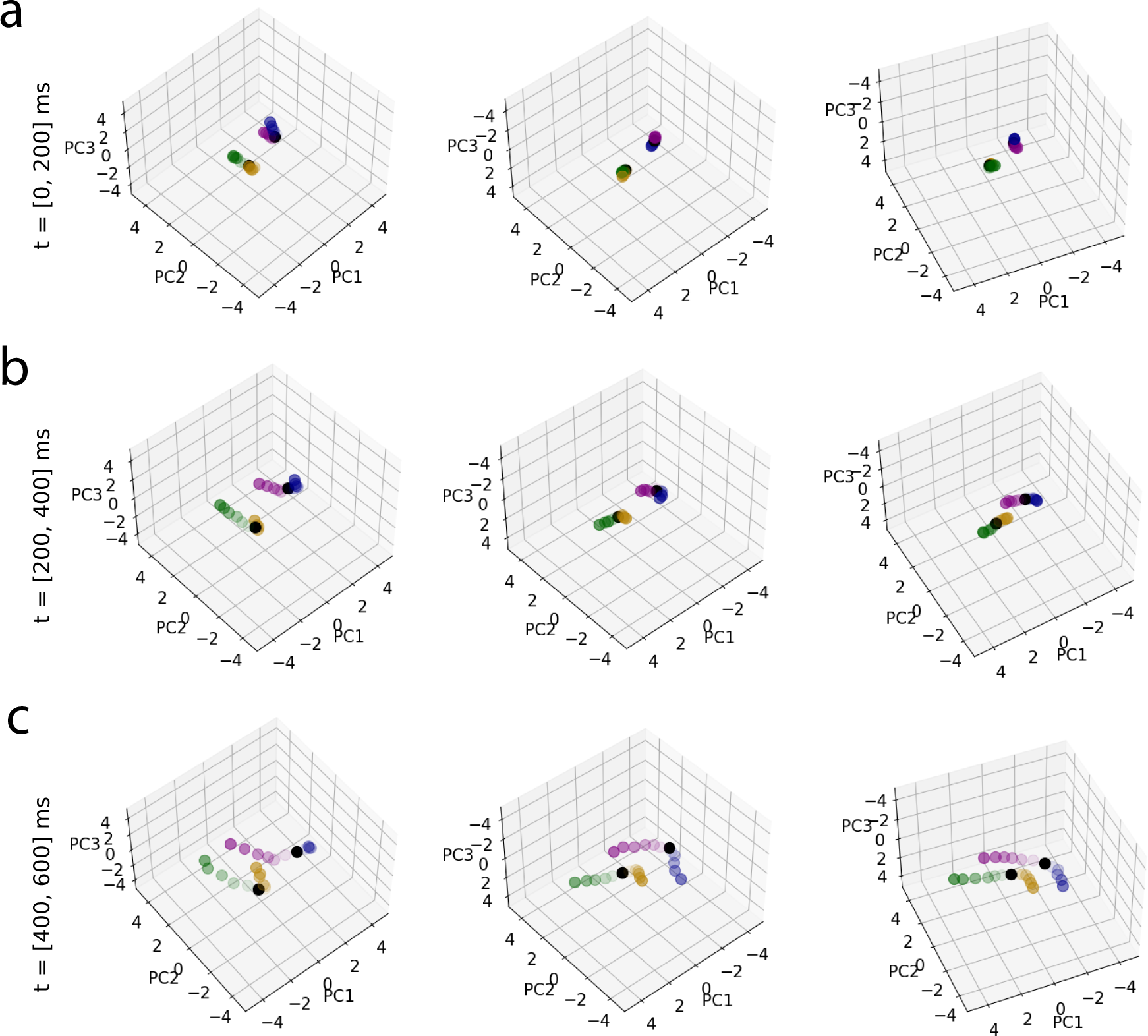
Context-dependent representational geometry in Monkey 1. lPFC population activity projected onto the top three principal components, shown for three representative recording sessions (columns) from monkey 1 in three intervals after stimulus onset (a-c). Conventions are similar to Fig. 3b.

**Figure S3:**
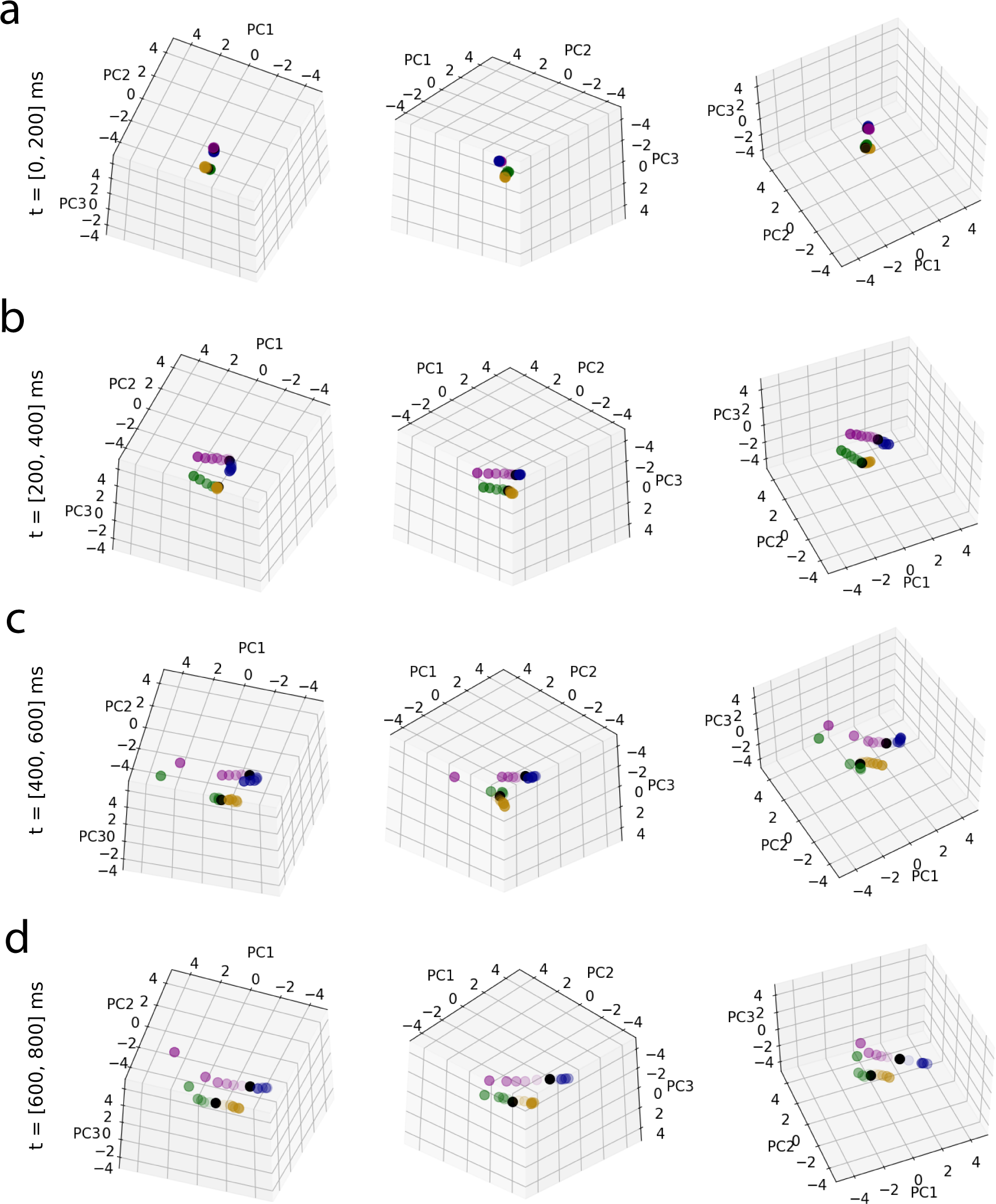
Context-dependent population geometry in Monkey 2.Same as Fig. S2, but showing three representa-tive sessions from monkey 2. Longer reaction times of this monkey allowed an additional analysis window: *t* = [600, 800] ms.

**Figure S4:**
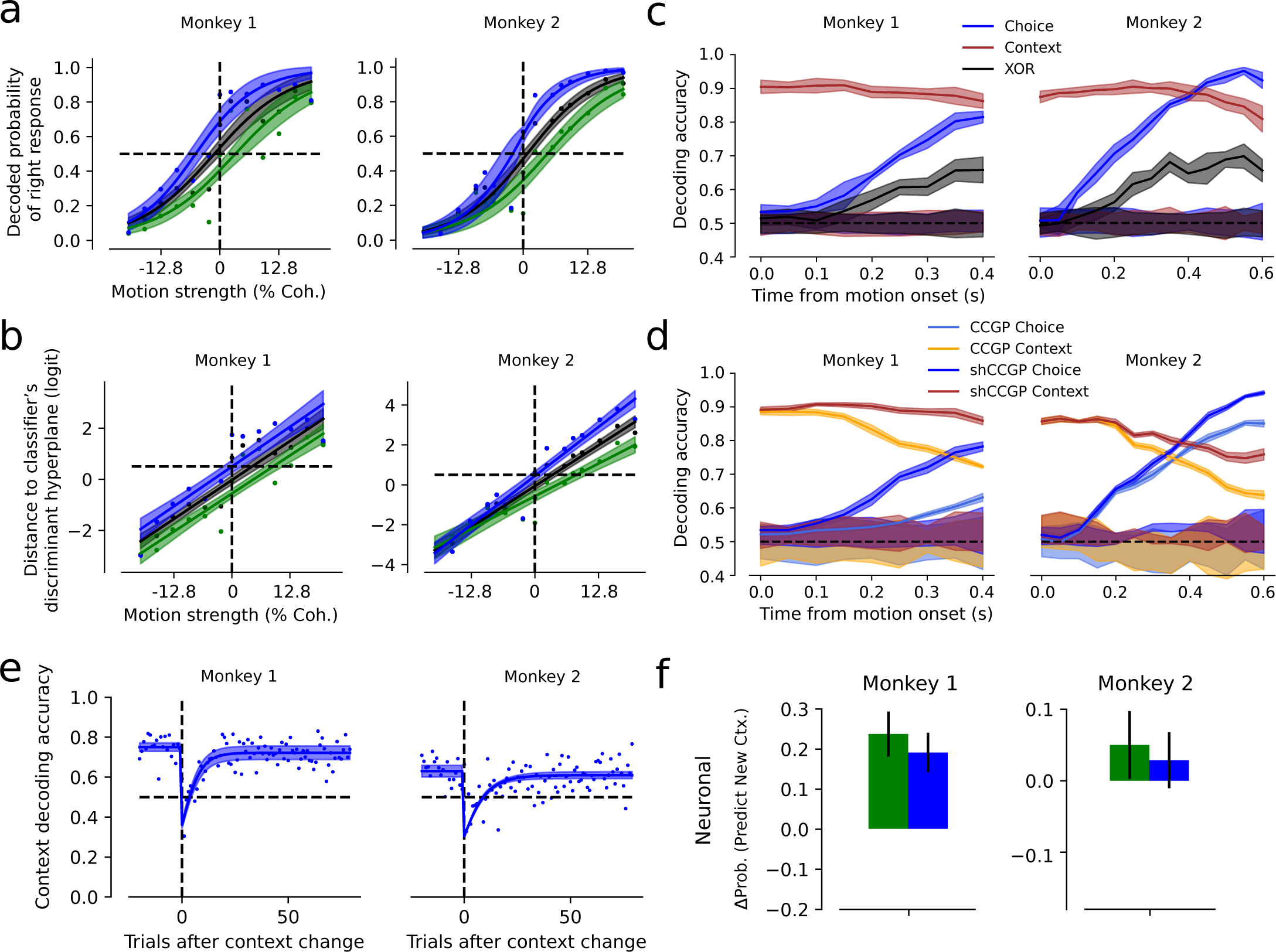
Neural representational geometry of individual monkeys. Same analysis as in Fig. 3, shown separately for monkey 1 and monkey 2.

**Figure S5:**
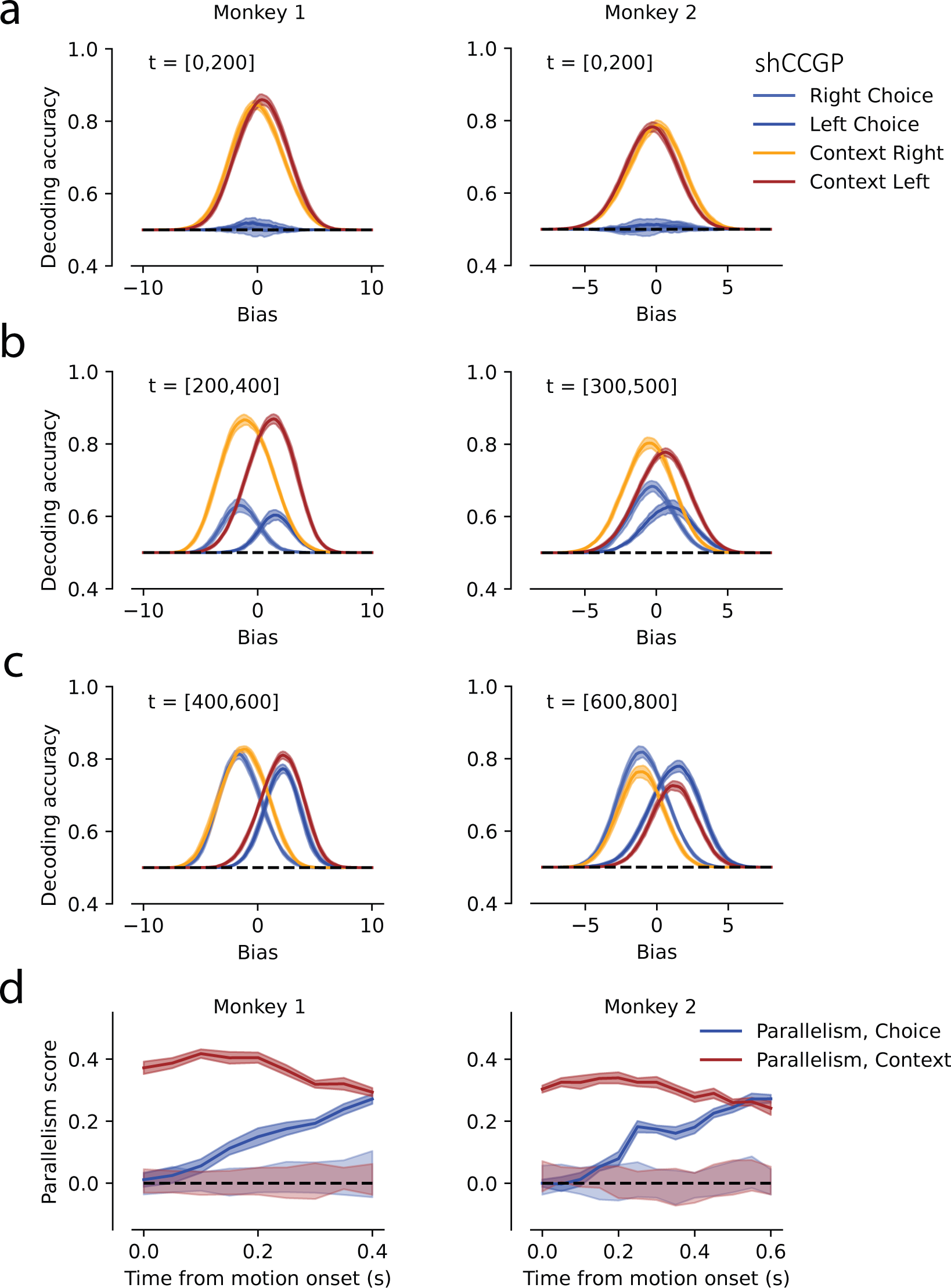
Shifted Cross-Condition Generalization Performance (shCCGP) for individual monkeys. (a) CCGP as a function of classifier bias offset (*ω*_0_) in an early post-stimulus window (*t* = [0, 200] ms) for decoding choice (blue) and reward context (orange) from lPFC population activity. Standard CCGP corresponds to bias = 0; shCCGP is defined as the maximum generalization performance across bias offsets. (b-c) Same analysis for later time windows. (d) Parallelism scores over time for choice and reward context.

**Figure S6:**
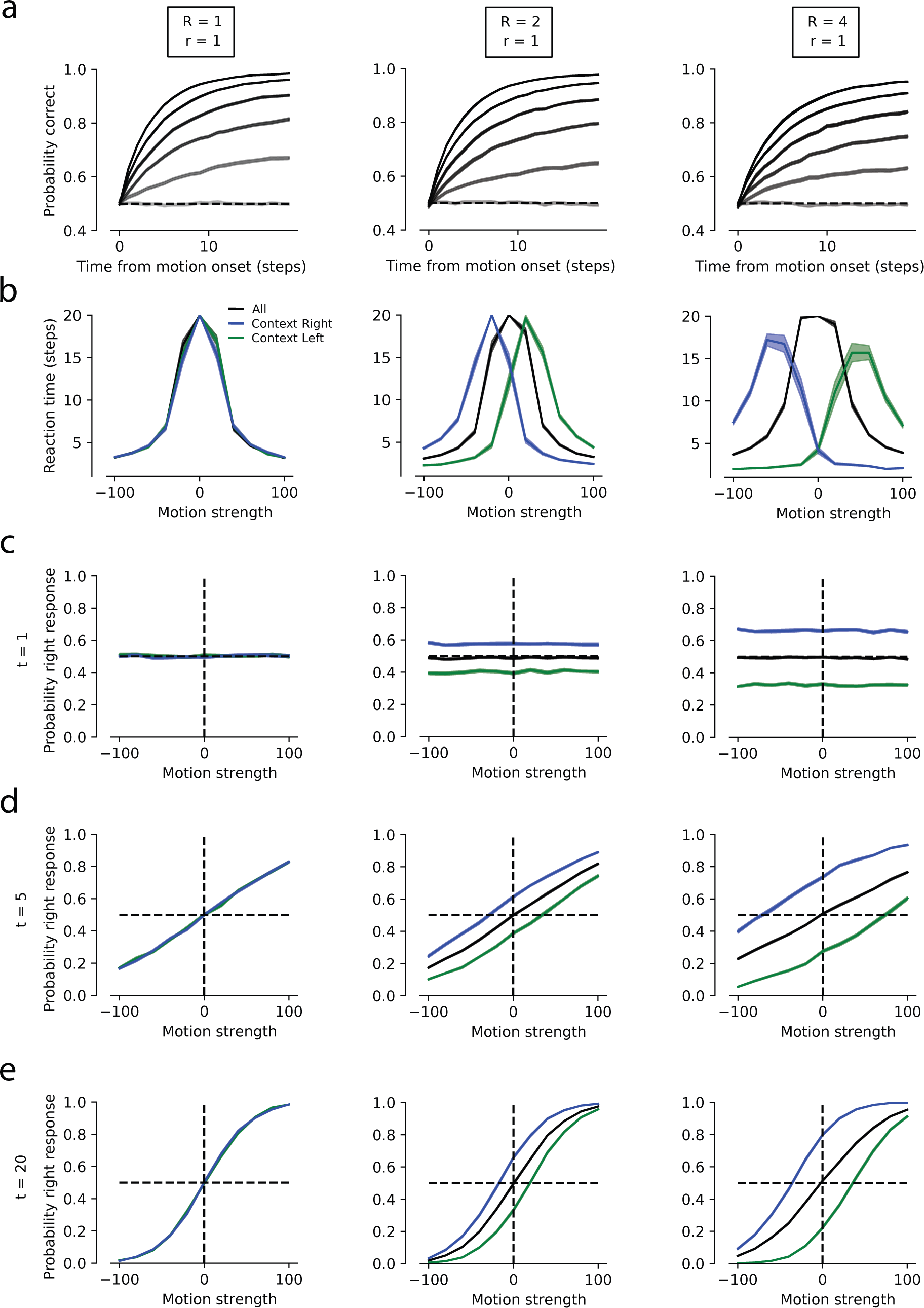
Increasing reward asymmetries enhanced behavioral bias in RNNs. (**a**) Probability of cor-rect responses as a function of stimulus presentation time for RNNs trained under different reward asymmetries ([*R, r*] ∈ {[1, 1], [2, 1], [4, 1]}). Gray scale indicates absolute motion strength, ranging from 0 (lightest gray) to 100 (darkest gray), evenly spaced. (**b**) Chronometric curve for left context (green), right context (blue), and all trials (black). (**c**-**e**) Psychometric curves at early (*t* = 1), intermediate (*t* = 5), and late (*t* = 20) time steps in simulated trials. Error bars denote s.e.m. across 50 networks.

**Figure S7:**
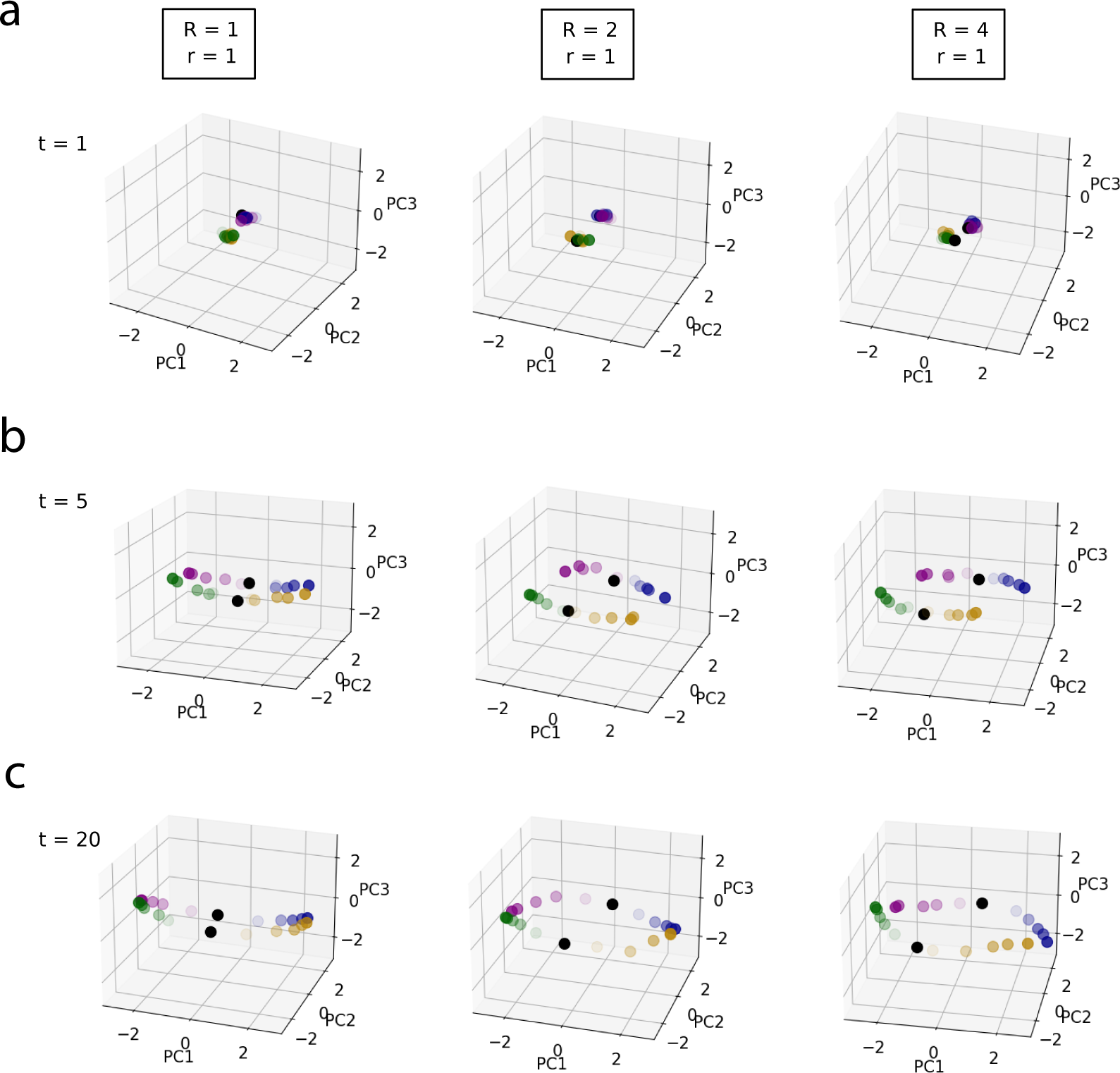
Reward asymmetry modulated RNN population geometry. PCA projection of the RNN activity under increasing reward asymmetry (columns) at early (a, *t* = 1), intermediate (b, *t* = 5), and late (c, *t* = 20) time steps in stimulated trials. Color conventions match Fig. 4c.

**Figure S8:**
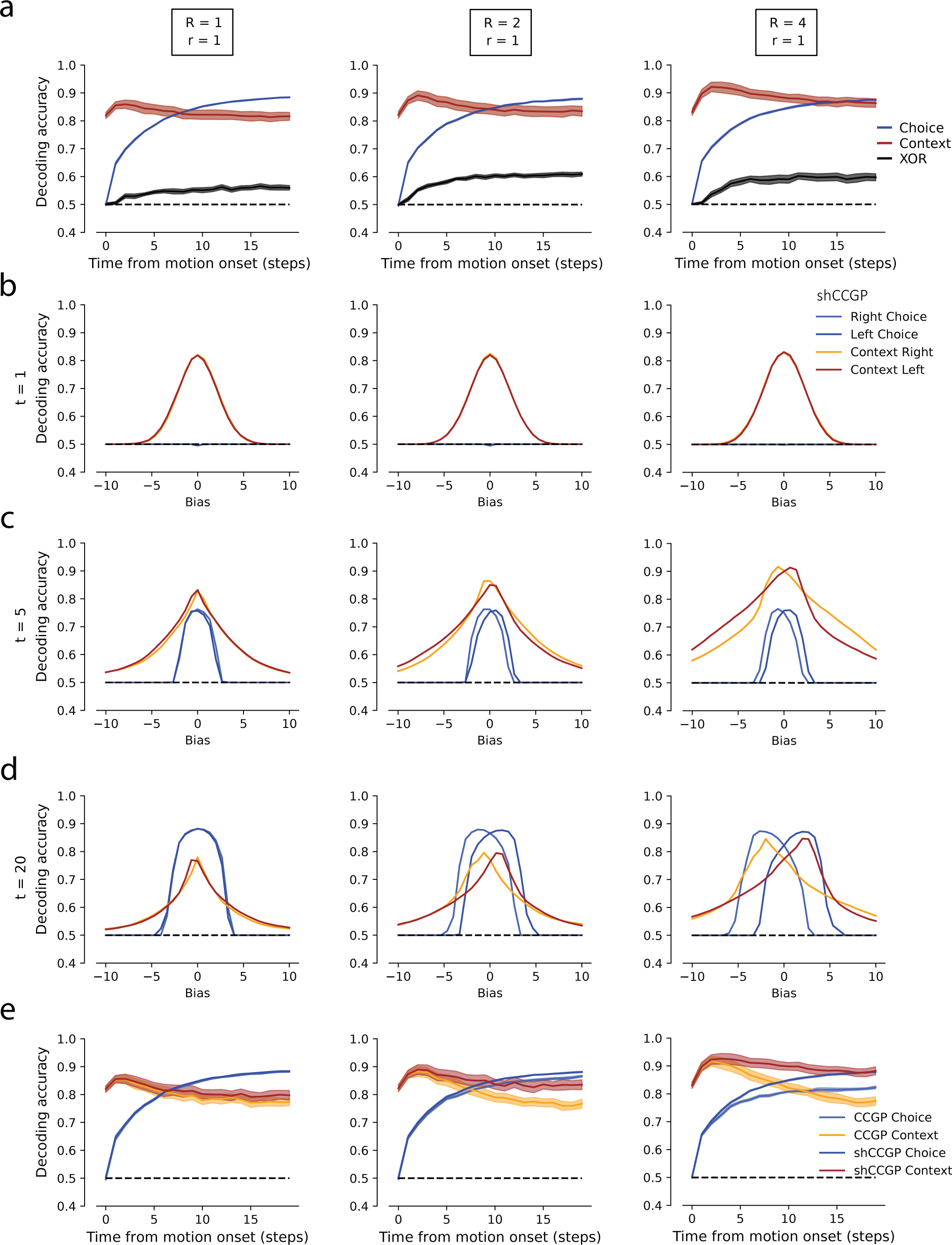
**Larger reward asymmetries enhanced representational shifts in RNNs**. (**a**) Decoding accuracy for choice (blue), context (red), and XOR between choice and context (black) as a function of time for different reward asymmetries (columns). (**b**-**d**) CCGP as a function of classifier bias offset at *t* = 1 (**b**), *t* = 5 (**c**), and *t* = 20 (**d**). (**e**) Standard CCGP (bias = 0) and shCCGP over time. Error bars indicate s.e.m. across 50 networks.

**Figure S9:**
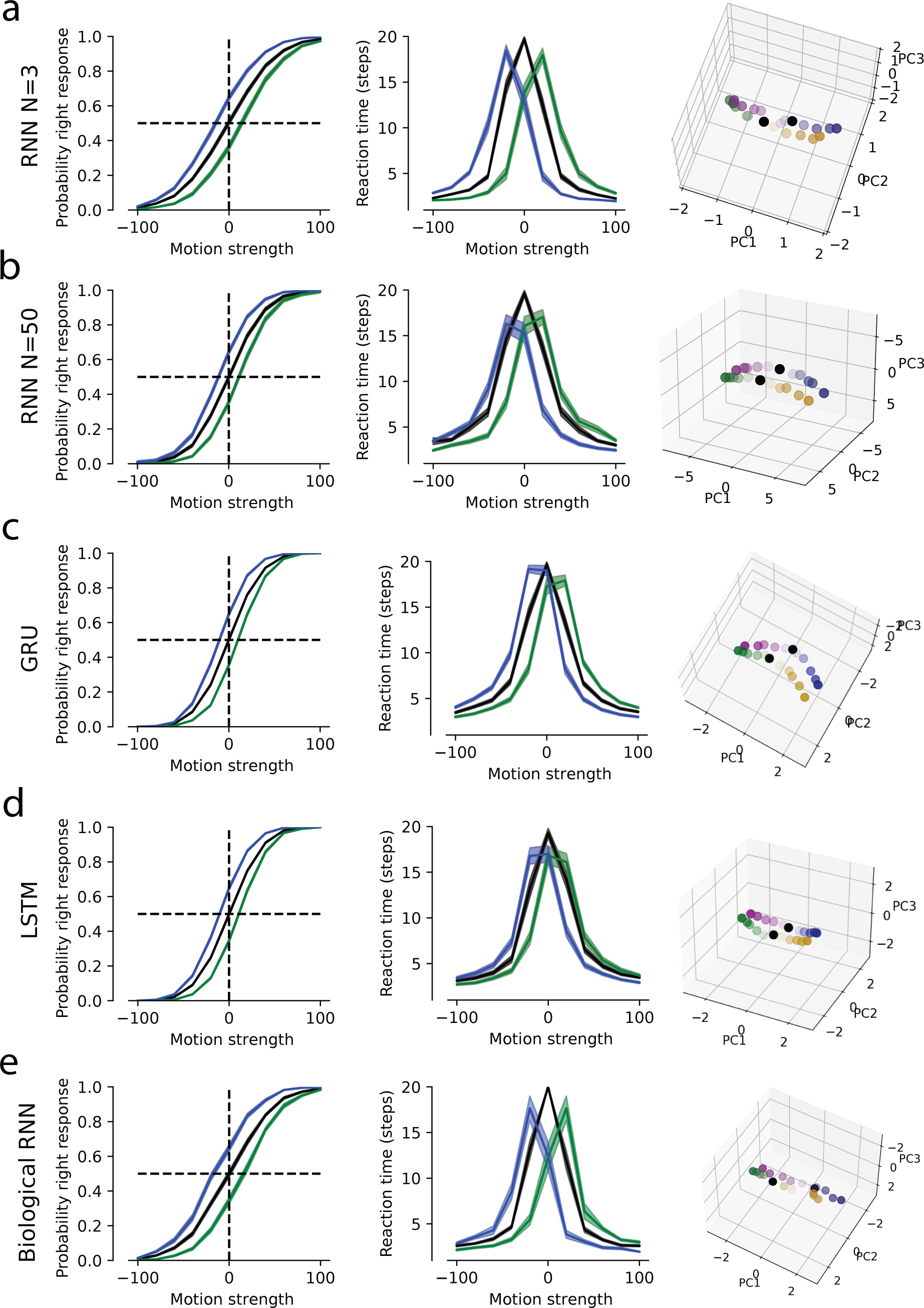
Diverse RNN architectures reproduced behavioral and representational geometry signatures. (a-b) Vanilla RNNs with three (a) and fifty (b) units reproduced the monkey’s psychometric and chronometric functions, as well as lPFC representational geometry (*t* = 5). (c) GRU architecture. (d) LSTM architecture. (e) Dale-constrained RNN (80% excitatory, 20% inhibitory units).

**Figure S10:**
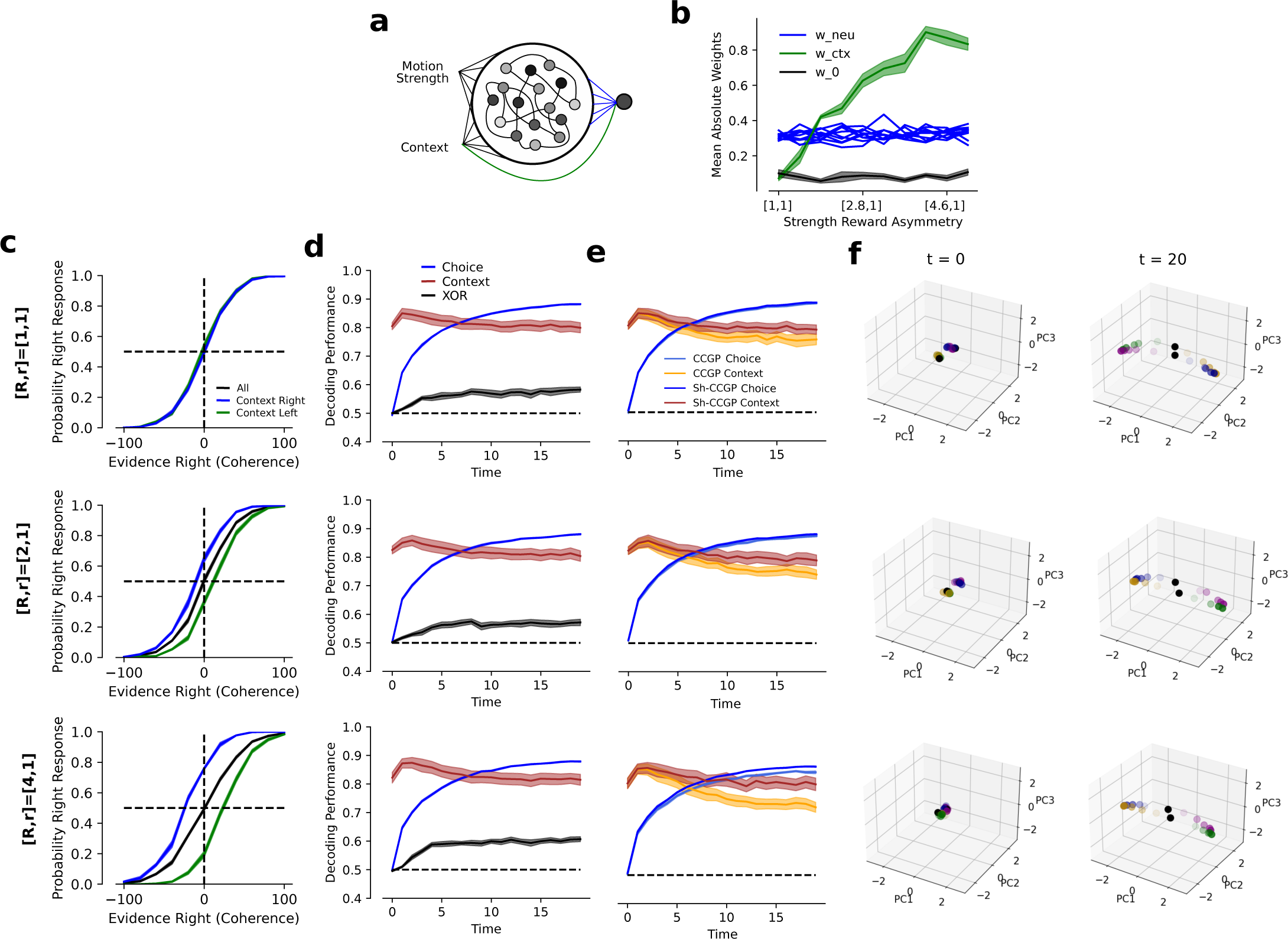
RNNs with explicit context signals sent to the readout unit shifted both decision variable manifold and the decision plane to generate the behavioral bias. (**a**) Modified RNN architecture in which the linear readout unit also received an explicit context signal. (**b**) Mean absolute weights of the readout unit as a function of reward asymmetry during training. The context weight (green, *w_ctx_*) increased with reward asymmetry, whereas readout weights from neurons (blue, *w_neu_*) and the intercept (gray, *w*_0_, decision plane) remained stable. The context weight quantified how much the decision plane shifted between the two contexts to create the behavioral bias. Error bars denote s.e.m. across 10 independent networks. (**c**-**f**) Psychometric function, decoding accuracy, CCGP/shCCGP, and PCA projections. Behavioral bias and decoding accuracies were identical to the RNNs without explicit context inputs to the readout unit, whereas representational shifts were smaller. Compare to Supplementary Figs. S6e, S7, and S8a,e. Error bars correspond to s.e.m. across 10 networks.

**Figure S11:**
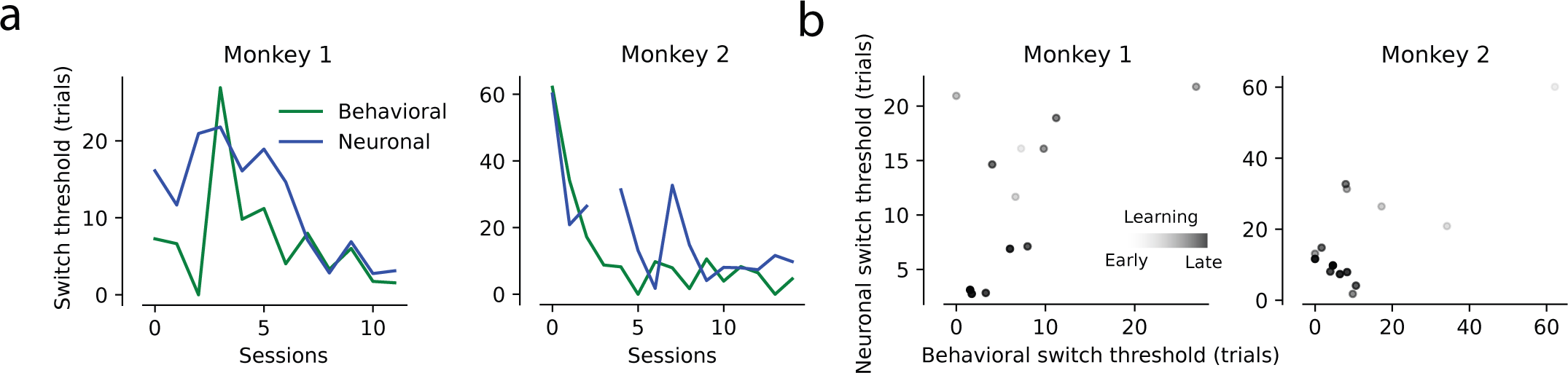
Both monkeys learned to infer reward context across training. (**a**) Behavioral (green) and neuronal (blue) thresholds across recording sessions for each monkey. Both monkeys learned to adapt their choices faster after a context switch across training. (**b**) Correlation between behavioral and neuronal switch thresholds across recording sessions for each monkey.

**Figure S12:**
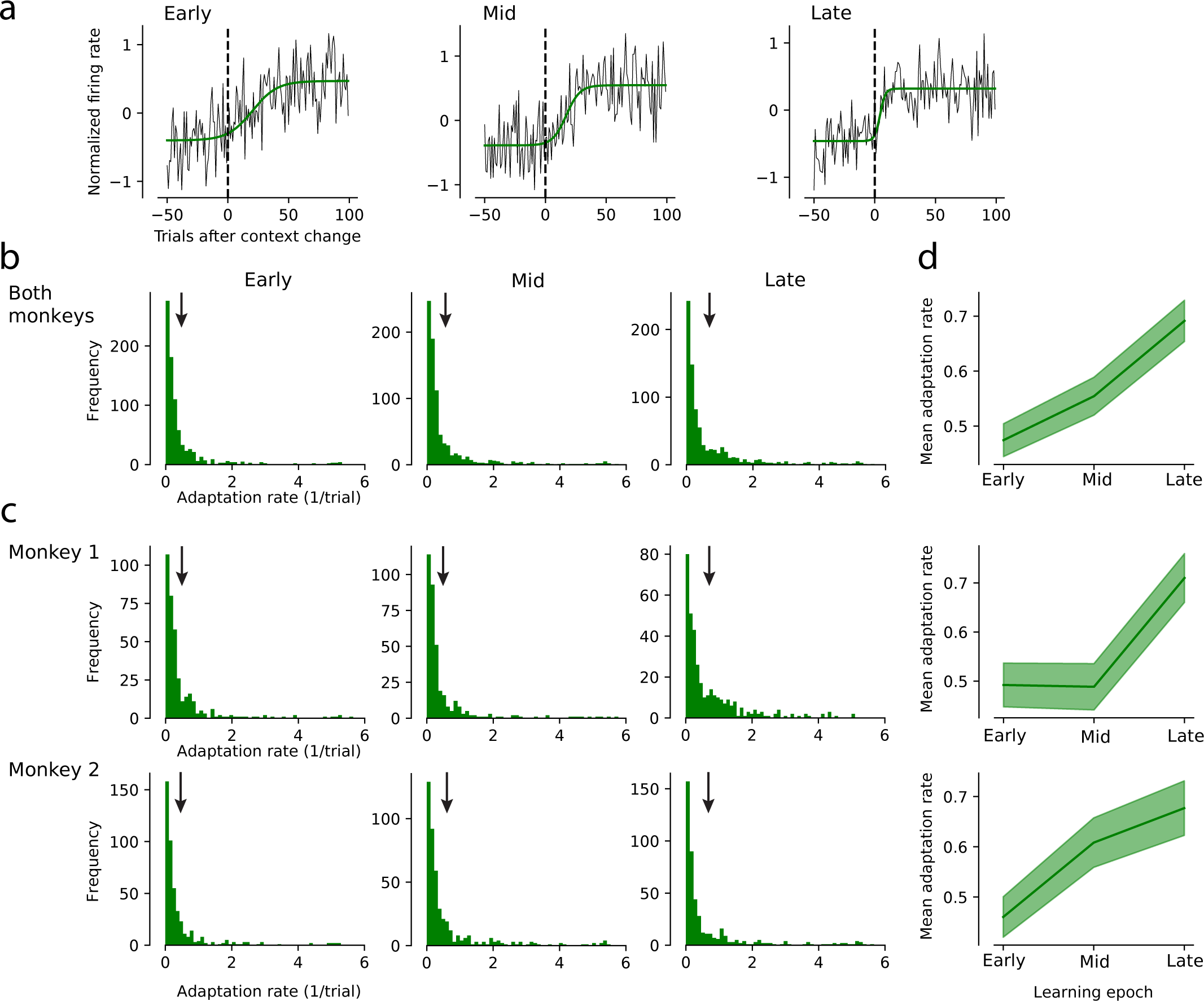
Individual unit activity dynamics reflect learning-dependent context inference. (a) Example units from monkey 1 showing faster post-switch adaptation from early (left) to mid (center), and late (right) stages of training. Firing rates at stimulus onset were fit with a sigmoid function (green line; Eq. 10; Monkey 1: *t* = [0, 200] ms; Monkey 2: *t* = [0, 300] ms). (b) Distribution of adaptation rates across learning epochs, quantified by the sigmoid slope parameter (*ω*_1_ in Eq. 10), pooled across both monkeys. Arrows indicate the mean for each training epoch (columns). (c) Same as b, shown separately for individual monkeys. (d) Changes of mean adaptation rate of individual units across learning epochs for both monkeys combined (top), monkey 1 (middle), and monkey 2 (bottom). Error bars denote s.e.m. across units.

**Figure S13:**
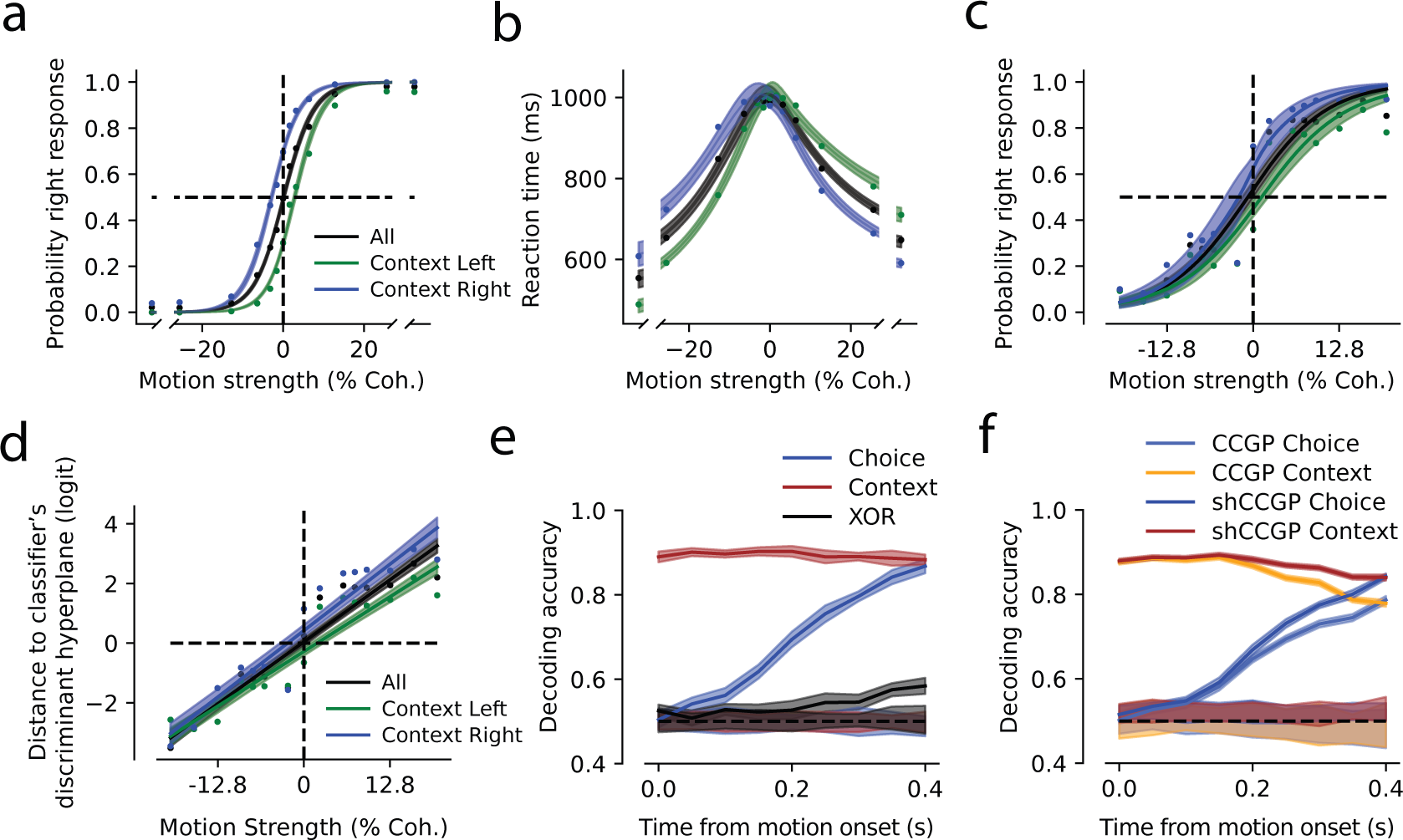
End-of-block choice bias and shifted lPFC representations are already present early in training. (**a**-**b**) Psychometric and chronometric curves from the first third of training sessions, computed from the late, within-block trials (last X trials per context block) to estimate asymptotic choice and RT bias. Plotting conventions match Fig. 2. (**c**-**f**) lPFC population representations from the same early sessions already exhibit context-dependent shifts between manifolds. Plotting conventions match Fig. 3. Together, these analyses show that steady-state behavioral bias and shifted representational geometry are present early in training, even though rapid post-switch adaptations (e.g., one-trial inference) develop later with training (see Fig. 5).

## References

[1] Purcell, B. A. & Kiani, R. Hierarchical decision processes that operate over distinct timescales underlie choice and changes in strategy. Proceedings of the National Academy of Sciences of the United States of America 113, E4531–4540 (2016).

[2] Okazawa, G. & Kiani, R. Neural mechanisms that make perceptual decisions flexible. Annual review of physiology 85, 191–215 (2023).

[3] Akrami, A., Kopec, C. D., Diamond, M. E. & Brody, C. D. Posterior parietal cortex represents sensory history and mediates its effects on behaviour. Nature 554, 368–372 (2018).

[4] Kamigaki, T., Fukushima, T. & Miyashita, Y. Cognitive set reconfiguration signaled by macaque posterior parietal neurons. Neuron 61, 941–951 (2009).

[5] Sarafyazd, M. & Jazayeri, M. Hierarchical reasoning by neural circuits in the frontal cortex. Science 364, eaav8911 (2019).

[6] Mansouri, F. A., Tanaka, K. & Buckley, M. J. Conflict-induced behavioural adjustment: a clue to the executive functions of the prefrontal cortex. Nature Reviews Neuroscience 10, 141–152 (2009).

[7] Shadlen, M. N. & Kiani, R. Decision making as a window on cognition. Neuron 80, 791–806 (2013).

[8] Levy, D. J. & Glimcher, P. W. The root of all value: a neural common currency for choice. Current opinion in neurobiology 22, 1027–1038 (2012).

[9] Lee, D., Seo, H. & Jung, M. W. Neural basis of reinforcement learning and decision making. Annual review of neuroscience 35, 287–308 (2012).

[10] Duan, C. A. et al. Collicular circuits for flexible sensorimotor routing. Nature neuroscience 24, 1110–1120 (2021).

[11] Hanks, T. D., Mazurek, M. E., Kiani, R., Hopp, E. & Shadlen, M. N. Elapsed decision time affects the weighting of prior probability in a perceptual decision task. Journal of Neuroscience 31, 6339–6352 (2011).

[12] Rao, V., DeAngelis, G. C. & Snyder, L. H. Neural correlates of prior expectations of motion in the lateral intraparietal and middle temporal areas. Journal of Neuroscience 32, 10063–10074 (2012).

[13] Mochol, G., Kiani, R. & Moreno-Bote, R. Prefrontal cortex represents heuristics that shape choice bias and its integration into future behavior. Current Biology 31, 1234–1244 (2021).

[14] Mansouri, F. A., Buckley, M. J. & Tanaka, K. Mnemonic function of the dorsolateral prefrontal cortex in conflict-induced behavioral adjustment. Science 318, 987–990 (2007).

[15] Hayden, B. Y., Pearson, J. M. & Platt, M. L. Neuronal basis of sequential foraging decisions in a patchy environment. Nature neuroscience 14, 933–939 (2011).

[16] Nogueira, R. et al. Lateral orbitofrontal cortex anticipates choices and integrates prior with current information. Nature Communications 8, 1–13 (2017).

[17] Findling, C. et al. Brain-wide representations of prior information in mouse decision-making. Nature 645, 192–200 (2025).

[18] Glimcher, P. W. Foundations of neuroeconomic analysis (Oxford University Press, 2010).

[19] Rangel, A., Camerer, C. & Montague, P. R. A framework for studying the neurobiology of value-based decision making. Nature reviews neuroscience 9, 545–556 (2008).

[20] Rorie, A. E., Gao, J., McClelland, J. L. & Newsome, W. T. Integration of sensory and reward information during perceptual decision-making in lateral intraparietal cortex (lip) of the macaque monkey. PloS one 5, e9308 (2010).

[21] Noorbaloochi, S., Sharon, D. & McClelland, J. L. Payoff information biases a fast guess process in perceptual decision making under deadline pressure: evidence from behavior, evoked potentials, and quantitative model comparison. Journal of Neuroscience 35, 10989–11011 (2015).

[22] Summerfield, C. & Koechlin, E. Economic value biases uncertain perceptual choices in the parietal and prefrontal cortices. Frontiers in human neuroscience 4, 208 (2010).

[23] Diederich, A. & Busemeyer, J. R. Modeling the effects of payoff on response bias in a perceptual discrimination task: Bound-change, drift-rate-change, or two-stage-processing hypothesis. Perception & Psychophysics 68, 194–207 (2006).

[24] Moran, R. Optimal decision making in heterogeneous and biased environments. Psychonomic bulletin & review 22, 38–53 (2015).

[25] Huang, Y., Hanks, T., Shadlen, M., Friesen, A. L. & Rao, R. P. How prior probability influences decision making: A unifying probabilistic model. Advances in neural information processing systems 25 (2012).

[26] Leon, M. I. & Shadlen, M. N. Effect of expected reward magnitude on the response of neurons in the dorsolateral prefrontal cortex of the macaque. Neuron 24, 415–425 (1999).

[27] Roitman, J. D. & Shadlen, M. N. Response of neurons in the lateral intraparietal area during a combined visual discrimination reaction time task. Journal of neuroscience 22, 9475–9489 (2002).

[28] Purcell, B. A. & Kiani, R. Neural Mechanisms of Post-error Adjustments of Decision Policy in Parietal Cortex. Neuron 89, 658–671 (2016).

[29] Waskom, M. L., Okazawa, G. & Kiani, R. Designing and interpreting psychophysical investigations of cognition. Neuron 104, 100–112 (2019).

[30] Fusi, S., Miller, E. K. & Rigotti, M. Why neurons mix: high dimensionality for higher cognition. Current opinion in neurobiology 37, 66–74 (2016).

[31] Kaufman, M. T. et al. The implications of categorical and category-free mixed selectivity on representational geometries. Current opinion in neurobiology 77, 102644 (2022).

[32] Ostojic, S. & Fusi, S. Computational role of structure in neural activity and connectivity. Trends in Cognitive Sciences 28, 677–690 (2024).

[33] Haxby, J. V. et al. A common, high-dimensional model of the representational space in human ventral temporal cortex. Neuron 72, 404–416 (2011).

[34] Guntupalli, J. S. et al. A model of representational spaces in human cortex. Cerebral cortex 26, 2919–2934 (2016).

[35] Bernardi, S. et al. The geometry of abstraction in the hippocampus and prefrontal cortex. Cell 183, 954–967 (2020).

[36] Fascianelli, V. et al. Neural representational geometries reflect behavioral differences in monkeys and recurrent neural networks. Nature Communications 15, 6479 (2024).

37. Courellis, H. S., et al. Abstract representations emerge in human hippocampal neurons during inference behavior. bioRxiv (2023).

[38] Higgins, I. et al. Unsupervised deep learning identifies semantic disentanglement in single inferotemporal face patch neurons. Nature communications 12, 6456 (2021).

[39] Nogueira, R., Rodgers, C. C., Bruno, R. M. & Fusi, S. The geometry of cortical representations of touch in rodents. Nature Neuroscience 1–12 (2023).

40. Boyle, L., Posani, L., Irfan, S., Siegelbaum, S. A. & Fusi, S. The geometry of hippocampal ca2 representations enables abstract coding of social familiarity and identity. bioRxiv (2022).

[41] Okazawa, G., Hatch, C. E., Mancoo, A., Machens, C. K. & Kiani, R. Representational geometry of perceptual decisions in the monkey parietal cortex. Cell 184, 3748–3761 (2021).

[42] Monsalve-Mercado, M. M., Stine, G. M., Shadlen, M. N. & Miller, K. D. The geometry of the neural state space of decisions. bioRxiv 2025–01 (2025).

[43] Rigotti, M. et al. The importance of mixed selectivity in complex cognitive tasks. Nature 497, 585–590 (2013).

[44] Wutz, A., Loonis, R., Roy, J. E., Donoghue, J. A. & Miller, E. K. Different levels of category abstraction by different dynamics in different prefrontal areas. Neuron 97, 716–726 (2018).

[45] Pagan, M. et al. Individual variability of neural computations underlying flexible decisions. Nature 639, 421–429 (2025).

[46] Link, S. W. The wave theory of difference and similarity (Routledge, 2020).

[47] Shadlen, M. N., Hanks, T. D., Churchland, A. K., Kiani, R. & Yang, T. The speed and accuracy of a simple perceptual decision: a mathematical primer. Bayesian brain: Probabilistic approaches to neural coding 209–237 (2006).

[48] Ratcliff, R. & McKoon, G. The diffusion decision model: theory and data for two-choice decision tasks. Neural computation 20, 873–922 (2008).

[49] Kim, J.-N. & Shadlen, M. N. Neural correlates of a decision in the dorsolateral prefrontal cortex of the macaque. Nature neuroscience 2, 176–185 (1999).

[50] Kiani, R., Cueva, C. J., Reppas, J. B. & Newsome, W. T. Dynamics of neural population responses in prefrontal cortex indicate changes of mind on single trials. Current Biology 24, 1542–1547 (2014).

[51] Mante, V., Sussillo, D., Shenoy, K. V. & Newsome, W. T. Context-dependent computation by recurrent dynamics in prefrontal cortex. nature 503, 78–84 (2013).

[52] Cueva, C. J. et al. Low-dimensional dynamics for working memory and time encoding. Proceedings of the National Academy of Sciences 117, 23021–23032 (2020).

[53] Yang, G. R. & Wang, X.-J. Artificial neural networks for neuroscientists: a primer. Neuron 107, 1048–1070 (2020).

[54] Ni, A. M., Ruff, D. A., Alberts, J. J., Symmonds, J. & Cohen, M. R. Learning and attention reveal a general relationship between population activity and behavior. Science 359, 463–465 (2018).

[55] Sanayei, M. et al. Perceptual learning of fine contrast discrimination changes neuronal tuning and population coding in macaque v4. Nature communications 9, 4238 (2018).

[56] Ruff, D. A. & Cohen, M. R. Simultaneous multi-area recordings suggest that attention improves performance by reshaping stimulus representations. Nature neuroscience 22, 1669–1676 (2019).

[57] Lam, N. H. et al. Prefrontal transthalamic uncertainty processing drives flexible switching. Nature 637, 127–136 (2025).

[58] Rikhye, R. V., Gilra, A. & Halassa, M. M. Thalamic regulation of switching between cortical representations enables cognitive flexibility. Nature neuroscience 21, 1753–1763 (2018).

[59] Brainard, D. H. The psychophysics toolbox. Spatial Vision 10, 433–6 (1997).

[60] Britten, K. H., Shadlen, M. N., Newsome, W. T. & Movshon, J. A. The analysis of visual motion: a comparison of neuronal and psychophysical performance. Journal of Neuroscience 12, 4745–4765 (1992).

[61] Kiani, R., Hanks, T. D. & Shadlen, M. N. Bounded integration in parietal cortex underlies decisions even when viewing duration is dictated by the environment. Journal of Neuroscience 28, 3017–3029 (2008).

[62] Trautmann, E. M. et al. Accurate estimation of neural population dynamics without spike sorting. Neuron 103, 292–308 (2019).

[63] Palmer, J., Huk, A. C. & Shadlen, M. N. The effect of stimulus strength on the speed and accuracy of a perceptual decision. Journal of vision 5, 1–1 (2005).

[64] Chen, G., Kang, B., Lindsey, J., Druckmann, S. & Li, N. Modularity and robustness of frontal cortical networks. Cell 184, 3717–3730 (2021).

[65] Bondy, A. G. et al. Coordinated cross-brain activity during accumulation of sensory evidence and decision commitment. bioRxiv 2024–08 (2024).

[66] Hochreiter, S. & Schmidhuber, J. Long short-term memory. Neural computation 9, 1735–1780 (1997).

67. Cho, K., et al. Learning phrase representations using rnn encoder-decoder for statistical machine translation. arXiv preprint arXiv:1406.1078 (2014).

